# Evolved and plastic gene expression in adaptation to a novel niche

**DOI:** 10.1101/2024.03.02.583104

**Authors:** Rachel A Steward, Jesús Ortega Giménez, Shruti Choudhary, Su Yi, Olivier Van Aken, Anna Runemark

## Abstract

Gene expression, resulting from complex regulatory interactions, plays an important role in adaptation and speciation. While gene expression historically has been studied in the context of reproductive isolation in speciation research, a role for evolved differences in gene expression in adaptation to novel niches is increasingly appreciated. How gene expression evolves and enables divergent ecological adaptation, and how changes in gene expression relate to genomic architecture and genetic divergence are pressing questions in understanding the processes of adaptation and ecological speciation. Further, how plasticity in gene expression can both contribute to and be affected by the process of ecological adaptation is a crucial component in understanding gene expression evolution. To address these questions, we investigate the role of evolved and plastic gene expression differences in adaptation leveraging an established host plant shift in the peacock fly *Tephritis conura*. Using a cross-fostering design where larvae feed on either natal or alternate host plants, we uncover extensive evolved differences in gene expression between the ecotypes, strikingly in genes associated with processing of host plant chemicals. We find limited evidence for plasticity, with some indications of higher plasticity in the ancestral ecotype where the expression of three gene coexpression modules is altered when larvae are cross-fostered to the derived host plant. Interestingly, we find an enrichment of differentially expressed genes within a large, ecotype-specific inversion in the *T. conura* genome. This finding adds to evidence that inversions are important for enabling diversification in the face of gene flow and underscores that effects on gene expression may be key to understanding the role of inversions.

## Introduction

Colonization of novel environments is a fundamental driver of biodiversity, often resulting in locally adapted, ecologically divergent populations, ecotypes or species (Rundle & Nosil, 2005). Uncovering the genomic changes underlying this process of adaptation is critical to understanding how the multifarious selection pressures of a novel environment can lead to ecological divergence and, ultimately, speciation (Wolfsberger et al., 2022). Over the past fifty years, it has become clear that sequence divergence at gene loci, especially within protein coding regions, is insufficient to explain major morphological, physiological, and behavioral divergence between species. Rather, mechanisms regulating expression of these genes are likely to account for biological and ecological divergence between populations, ecotypes and species (Barrier et al., 2001; Bawin et al., 2024; Carroll, 2000; King & Wilson, 1975; Rivas et al., 2018; Wittkopp & Kalay, 2012). These mechanisms include intricate combinations of coding and noncoding sequences that depend on genome architecture, structural variation, and epistatic interactions between regions of the genome (Berdan et al., 2023; Diehl et al., 2020; Drinnenberg et al., 2019; Emerson & Li, 2010; Fuller et al., 2016; Tirosh et al., 2010; Wittkopp & Kalay, 2012). Because small changes in coding or epigenetic regulatory mechanisms can have large, cascading effects in coexpressed gene networks, significant expression differences can evolve between lineages adapting to different niches even over short evolutionary timescales, with large consequences for phenotypes under selection in new environments. Evolved differences in gene expression not only create important barriers to recolonization of alternative niches but may also lead to regulatory changes that limit opportunities for gene flow between divergent lineages. In particular, hybrids between locally adapted or speciating populations may be unable to express the combination of genes necessary to succeed in either niche. Thus, understanding the evolution of expression during colonization of and adaptation to novel niches can inform our understanding of ecological speciation.

Gene expression is a particularly compelling source of phenotypic variation because, in part due to diverse regulatory mechanisms, it has the potential to respond plastically to different environments (Rivera et al., 2021; Schlichting & Pigliucci, 1993; Schlichting & Smith, 2002). Both evolved and plastic expression differences have been found to play a role in local adaptation and ecological divergence in novel environments (Albecker et al., 2021; Ballinger et al., 2023; Campbell-Staton et al., 2017; Ghalambor et al., 2015; Mack et al., 2018; Schoville et al., 2012; Y.-X. Xu et al., 2023). However, the relative importance of evolved and plastic expression differences along the speciation continuum is unclear. Both adaptive and non-adaptive plasticity have the potential to promote population persistence upon colonization of a novel niche (Ghalambor et al., 2007; Pavey et al., 2010), and might even accelerate adaptive evolution in response to the novel environment (Levis & Pfennig, 2020; Ng & Kinjo, 2023).

Ancestral plasticity is an important source of phenotypic variation that allows for colonization and ultimately divergence on a novel niche, as described in the flexible stem (Gibert, 2017; West-Eberhard, 2003) and oscillation (Janz & Nylin, 2008) hypotheses. One prediction of the flexible stem hypothesis is that strong selection of adaptive gene expression in the novel niche will ultimately lead to genetic accommodation or assimilation, a decrease or loss of ancestral plasticity in the derived lineage (Gibert, 2017). This has been observed in both natural and experimentally evolved populations that are adapted to divergent environments (Brennan et al., 2022; Kelly, 2019; Wood et al., 2023). In contrast, Celorio-Mancera et al. (2023) found that even specialized species of *Polygonia* butterflies retained gene expression modules associated with feeding on a diverse range of host plants. Further, in a reanalysis of published data, Chen and Zhang (2023) found that genetic assimilation was a rare outcome for ancestrally plastic genes. Thus, one major question about the evolution of gene expression during divergence on a novel niche is whether populations in the ancestral niche exhibit greater adaptive transcriptional plasticity than populations exploiting the derived niche.

Host plant shifts provide excellent study systems for examining the role of gene expression in adaptation to novel environments. Colonization of novel host plants and subsequent divergence, or even speciation, is especially common among herbivorous insects (Drès & Mallet, 2002; Hernández-Hernández et al., 2021; Jaenike, 1990). Hernandez-Hernandez et al. (2021) found that over 70% of reviewed sister species pairs were ecologically divergent, especially in their interactions with host plants. Herbivorous insects are often highly dependent on their host plants throughout their juvenile and adult lives (Ehrlich & Raven, 1964; Janz, 2011; Jermy, 1984), and host plants represent complex, multidimensional niches with multifarious selection pressures affecting insect fitness. Host shifts are also interesting because they are hypothesized to involve an initial period of niche polymorphism or plasticity before ecological specialization and speciation (Drès & Mallet, 2002; Janz & Nylin, 2008; Nyman et al., 2010). In fact, major patterns of herbivorous insect diversification can be linked to host use variability, rather than abrupt switches between host plants (Braga et al., 2018), suggesting an important role for phenotypic plasticity in adaptation and divergence (i.e., the oscillation hypothesis c.f. Janz & Nylin, 2008).

In addition to this macroevolutionary role, contemporary plasticity, especially transcriptional plasticity, could buffer against maladaptive host use, facilitate reacquisition of historical hosts or lead to colonization of additional host plants. Transcriptional divergence between insect species using different host plants or responding to dietary manipulations has been extensively studied (Birnbaum & Abbot, 2020). Counter to the predictions of the oscillation hypothesis, many expression differences between closely related species feeding on different host plants are constitutive, fixed differences, not transcriptionally plastic (Eyres et al., 2016; Orsucci et al., 2018; Silva-Brandão et al., 2017). This is consistent with evidence that there tend to be more *cis*-regulatory variants between species than within species, suggesting that the accumulation of targeted, regulatory elements is important to the process of divergence and speciation (Coolon et al., 2014; Emerson & Li, 2010; Hill et al., 2021; Tirosh et al., 2010) Thus, despite a hypothesized role for plasticity in colonization of and adaptation to novel hosts, few studies directly assess evolved versus plastic transcriptional differences in ongoing ecological speciation (Birnbaum & Abbot, 2020) how expression differences relate to underlying genomic architecture, and whether plasticity is likely to differ between ancestral and derived niches.

Here, we investigate the roles of gene expression divergence and plasticity underlying ongoing adaptation by the Peacock fly, *Tephritis conura*, to a novel host plant (Figure 1A). In continental Europe, *T. conura* populations specialize on *Cirsium heterophyllum* (Figure 1B) or *C. oleraceum* (Figure 1C) thistles, laying eggs and spending larval and pupal stages within a single thistle bud (Romstöck-Völkl, 1997). Broad geographical sampling suggests that *C. heterophyllum* was the ancestral host, and *C. oleraceum* has been colonized evolutionarily recently (est. 0.1-0.5MYA; Nilsson et al., 2023). Flies developing in the wrong host experience the highest mortality in the larval stage (Diegisser et al., 2008; Nilsson et al., In prep). As an incipient speciation event where there is still considerable gene flow in parapatric and sympatric contact zones (Nilsson et al., 2023), this system fills an important gap in research on ecological speciation (Anderson et al., 2023; Byers et al., 2017). *Cirsium heterophyllum* and *C. oleraceum* specialist lineages (CH and CO, respectively, Figure 1D) have a linked genomic basis of host use and reproductive isolation between ecotypes in the form of a large putative inversion on the ancestral dipteran X chromosome (Nilsson et al., 2023). However, the extent to which expression divergence and plasticity contribute to divergent host use, and whether these expression differences are associated with the putative inversion, are unknown.

**Figure 1.**
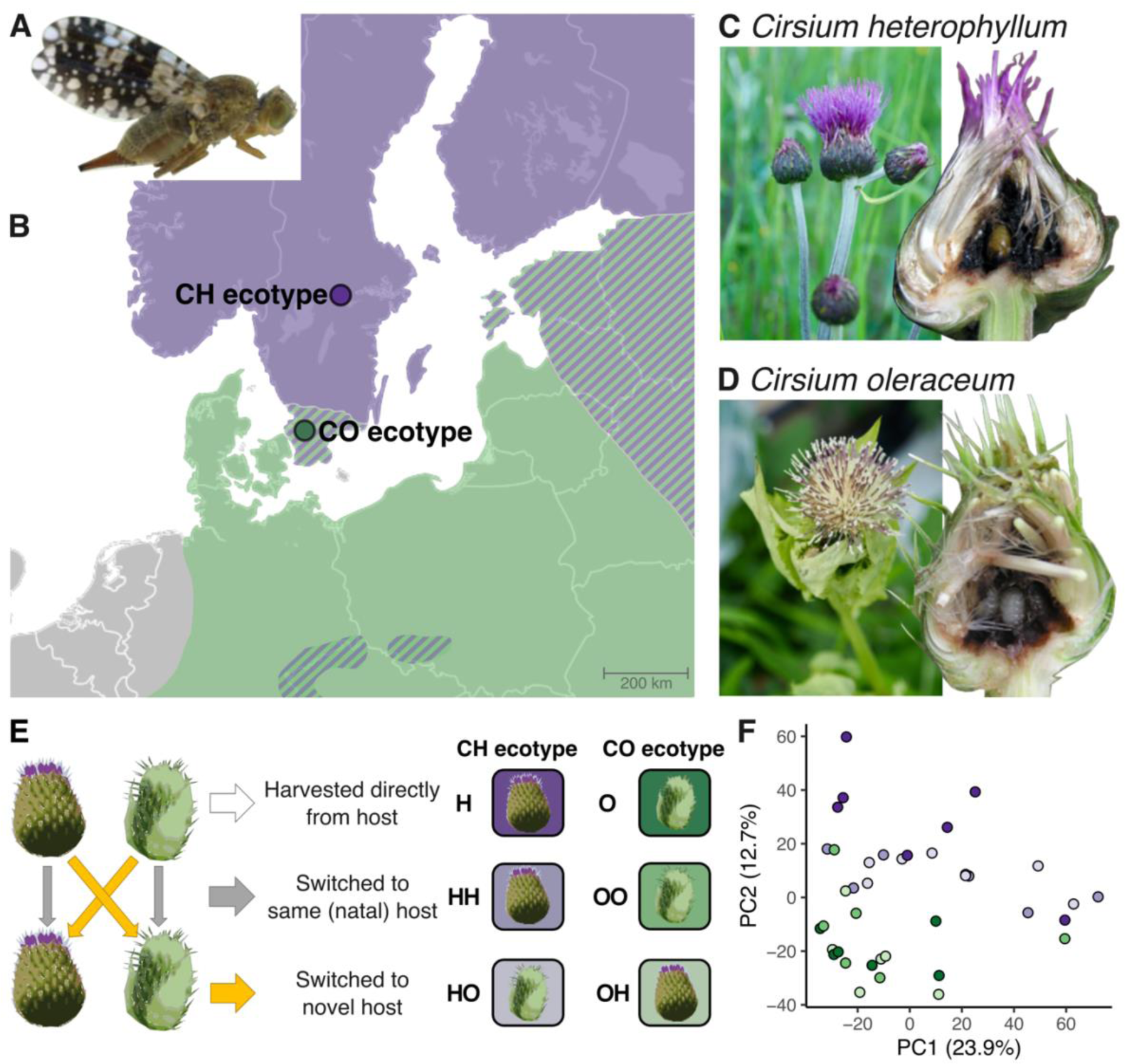
*Tephritis conura* host plants, sampling sites and cross-fostering design. (A) adult female *T. conura*. Image © K.J. Nilsson, used with permission. (B) *Cirsium heterophyllum* (CH) thistle buds, bisected bud showing fly pupae. Images © K.J. Nilsson, T. Diegisser, used with permission. (C) *C. oleraceum* (CO) thistle buds, bisected bud showing fly larvae. Images © K.J. Nilsson, T. Diegisser, used with permission. (D) Host plant distribution and sampling locations of buds containing CH and CO ecotypes; purple = allopatric *C. heterophyllum*, green = allopatric *C. oleraceum*, and striped = sympatric regions. (E) Cross fostering design and resulting treatments. (F) Principal component analysis of the 500 most variable genes expressed by third instar larvae. Points have the same colors as the squares in the cross-fostering design.

We specifically test the extent to which expression differs between *T. conura* ecotypes to understand the role of gene expression divergence in niche colonization and incipient speciation. Using a cross-fostering design (Figure 1D), we further test whether *T. conura* flies can plastically adjust gene expression in response to feeding on the alternate thistle host and whether the ancestral or derived ecotypes differ in the extent of their expression plasticity. Finally, we address how genomic architecture and divergent selection translate into constitutive and plastic gene expression differences between *T. conura* ecotypes. We find extensive expression differences between the two ecotypes feeding on their natal hosts, but limited evidence for gene expression plasticity when switched to the alternate host. However, a coexpression network approach suggests that some coexpressed gene modules can be plastically expressed in the ancestral ecotype. Finally, we find that constitutive expression differences between ecotypes were especially likely to occur in genes within the highly divergent putative inversion, suggesting a concerted role for genetic architecture and gene expression evolution in *T. conura* speciation.

## Methods

### Sample collection

We tested for differences in gene expression between divergent, host-associated ecotypes of the peacock fly *Tephritis conura* (Figure 1A). In northern parts of Scandinavia, populations of this fly specialize on the melancholy thistle, *Cirsium heterophyllum* (Figure 1C), while in southern parts of the range, flies lay their eggs on the cabbage thistle, *C. oleraceum* (Figure 1D), referred to as CH and CO flies, respectively. These lineages are often referred to as host races, however, here we use the broader term ecotype. Although the ranges overlap broadly and there are few morphological differences between these ecotypes (Nilsson et al., 2022), ecotypes show considerable genomic divergence and reduced gene flow in contact zones both east and west of the Baltic Sea (Nilsson et al., 2023; Figure 1B). Fitness costs of using the wrong host are highest in the larval stage (Diegisser et al., 2008), and we therefore focus on gene expression differences in third instar larvae feeding on natal or novel host plants. Larvae used in this experiment were sourced from a CH specialist population in central Sweden (59.63N, 14.58E) where *C. heterophyllum* is allopatric, and a CO population in southern Sweden (55.90N, 13.41E) where the host plants ranges are broadly sympatric but where *C. heterophyllum* is locally absent. Cross-fostering experiments were performed on *C. heterophyllum* and *C. oleraceum* plants in common garden facilities at Lund University, Lund, Sweden (Supplementary Methods).

To explore gene expression differences between the two ecotypes, third instar larvae were extracted from infested buds of each host plant and preserved for RNA extraction. We used relative sizes to estimate larval instar. The flies sampled from the original host plants form two control groups: H and O (Figure 1E). For the cross-fostering treatments, third instar larvae were extracted from infested buds and assigned to alternative host plants. Half of the larvae were switched into uninfested buds from the novel host plant, while the remaining larvae were switched to new buds of the natal host plant. This second group was included to control for gene expression differences resulting from the stress of moving larvae between thistle buds. After 6 hours feeding in the new bud, larvae were again extracted and preserved for RNA extraction. We confirmed that larvae fed on the new host by checking the inside of the bud for feeding damage and/or frass. The result was four cross-fostering treatments: CH larvae switched to *C. heterophyllum* buds (HH), CH larvae switched to *C. oleraceum* buds (HO), CO larvae switched to *C. oleraceum* buds (OO), CO larvae switched to *C. heterophyllum* buds (OH) (Figure 1E; Table S1; Supplementary methods).

### RNA extraction, library prep and sequencing

Extractions were performed using Sigma Aldrich’s Plant RNA kit, which we have found performs best with these flies. After sample quality testing, Illumina TruSeq Stranded mRNA libraries were prepared by SciLifeLab (Stockholm, Sweden). Mature transcripts were sorted using Poly-A selection. Sequencing was performed by SciLife Lab using theNovaSeq6000 (NovaSeq Control Software 1.7.5/RTA v3.4.4; ‘NovaSeqXp’ workflow in ‘S4’ mode flowcell). Raw read quality was assessed using FastQC (v. 0.11.9; Babraham Bioinformatics, https://www.bioinformatics.babraham.ac.uk/projects/fastqc/) and then adapters and low quality reads were trimmed using TrimGalore (v. 0.6.1; Ewels et al., 2016), which is a wrapper for CutAdapt (Martin, 2011). We specified a quality threshold of 10, excluding very low quality bases but is not likely to negatively affect mapping and differential expression results (Williams et al. 2016). Following trimming, reads were again assessed with FastQC and MultiQC (v.1.12; Ewels et al., 2016). On average, trimmed files contained 25.3 +/-0.8 M reads (Table S2).

### Read mapping and transcript quantification

To quantify gene expression, trimmed RNAseq reads were aligned against the library of coding transcripts from our in-house annotation (Supplementary Methods; Figure S1) using Salmon (v. 20180926; Patro et al., 2017). We ran Salmon using default parameters, specifying FR strandedness (ISF flag) and the –gcBias flag, as insects tend to have lower GC content than many model systems. Transcript counts (Transcripts Per Million) were concatenated into a single TXImport object for downstream gene-level analyses in the statistical platform R (v. 4.2.3;(R Core Team, 2023) using an inhouse R script (supplementary materials, salmon2dds.R) and the packages GenomicFeatures (v.1.52.2; Lawrence et al., 2013), tximport (v.1.28.0; Soneson et al., 2016), supported by the tidyverse (Wickham et al., 2019). One of the samples (P18653_176, OO treatment) had very poor alignment and quantification rates and was excluded. Otherwise, the proportion of trimmed reads aligned and quantified against transcripts ranged between 30-55% and did not differ between CH and CO ecotypes (Supplementary Figure S2).

### Differential expression analysis

To identify genes that were differentially expressed depending on ecotype and cross-fostering, differential gene expression analysis was performed using the DESeq2 package (v.1.36.0; Love et al., 2014) in R. Transcript abundance estimates were converted into gene abundance estimates, which were pre-filtered, keeping only those genes with at least 5 reads in a minimum of 6 samples, leaving 11701 expressed genes. Read counts were normalized for visualization and unsupervised clustering of gene expression using the ‘regularized log’ transformation implemented in DESeq2. To uncover major patterns of gene expression, a principal component analysis (stats::prcomp(); v.4.2.3; R Core Team, 2019) was performed on the 500 genes with the most variable expression after normalization.

We tested for pairwise differential expression among treatments (see Table S2 for full list of comparisons) by first fitting a model to the filtered set of genes with Treatment as the predictor (aka ‘design = ∼Treatment’ in DESeq2 nomenclature). We increased minReplicatesForReplace to 8 (default = 7), to make sure that outliers were refit with group averages for all contrasts, reducing the chance of false positives. We calculated log fold changes and assessed significant differences between treatment pairs by applying the adaptive Student’s t prior shrinkage estimator (apeglm). We used the shrinkage estimator to calculate the s-value (false sign rate, FSR; Stephens, 2017) instead of the adjusted p-value (false discovery rate, FDR). Rather than evaluating whether the difference between two groups is zero, the FSR tests the probability that the sign of the effect is likely to be true (Stephens, 2017). One motivation for using the s-value for our data was that we found considerable within group variation in our samples. After adjusting log fold changes for this variability using the DESeq2 function lfcshrink(), we still found highly significant adjusted p-values for very small log fold changes (e.g., LFC = 0.001 or a change of 1.00069%). While very small changes in gene expression can have phenotypic effects, we preferred to focus on those genes for which we were confident in the sign of the effect. As recommended, we used a smaller significance threshold than generally used for an FDR. We found that a threshold of s < 0.001 identified similarly sized gene sets as an FDR of 0.05 (https://support.bioconductor.org/p/133091/; Stephens, 2017; Zhu et al., 2019). Pairwise comparisons were visualized with volcano plots (-log10(s-value) against log2 fold change) and scatter plots comparing estimated log2 fold changes between specific sets of contrasts.

### Weighted gene coexpression network analysis

To further explore ecotype-specific gene expression and the extent to which the ecotypes can plastically adjust gene expression when feeding on different hosts, we identified groups or modules of genes with similar expression across samples using a weighted gene coexpression network analysis with WGCNA (v.1.72-5; Langfelder & Horvath, 2008, 2012) on the regularized log-transformed expression estimates. The soft thresholding power was selected as the first scale free topology fit index to exceed 0.8, using signed correlations (Langfelder & Horvath, 2008). We specified a signed network using biweight midcorrelation (bicor), required a minimum module size of 30, and merged any modules that were over 70% related. The biweight midcorrelation is more robust to outliers than is a standard Pearson correlation (Langfelder & Horvath, 2008).. Both the minimum module size and the merge-cut-height were chosen to reduce the number of very small modules. A module membership threshold of 0.6 was used to exclude genes with a poor fit in any given module. Genes with high membership in a given module are likely to represent highly interconnected “hub” genes (Langfelder & Horvath, 2008).

To uncover the factors affecting module expression, we correlated the standardized expression estimates for each module, called eigengenes, with each of 5 binary variables: ecotype, stress, reciprocal plasticity, CH plasticity and CO plasticity (Figure S3, Tables S1). We calculated Pearson’s correlation coefficients and student’s asymptotic p-values, which were corrected for multiple comparisons across modules using the Benjamini-Hochberg method (n = 21 tests). We visualized these correlations across modules using heatmaps (ComplexHeatmap v. 2.12.1; Gu, 2022; Gu et al., 2016).

### Functional enrichment of gene sets

To identify gene functions overrepresented among genes that were differentially expressed between ecotypes or in response to cross fostering treatment, functional enrichment was performed with TopGO (v.2.48.0; Alexa & Rahnenfuhrer, 2016) using a functional annotation made with EggNOG mapper (http://eggnog-mapper.embl.de/; v2 (Cantalapiedra et al., 2021; Huerta-Cepas et al., 2019; Supplementary methods). We required a minimum of 20 genes to perform gene set enrichment analysis (GSEA) and excluded GO terms that were associated with fewer than five genes. GSEA tested for overrepresentation of GO terms using one-sided Fisher’s exact tests (parent-child algorithm). A threshold of p < 0.05 was set to identify significantly enriched GO terms describing biological processes and molecular functions.

### Population differentiation, divergence, and signatures of selection

We next aimed to test the extent to which expression differences between ecotypes were associated with regions of genomic divergence between the two ecotypes, specifically a putative inversion segregating between the two ecotypes (Nilsson et al., 2023). We used site allele frequency files from Nilsson et al. (2023) to calculate Fst and dxy between the CH and CO source populations and to estimate nucleotide diversity (π) and Tajima’s D over 50kb non-overlapping windows throughout the 1.9G *T. conura* genome. The two-dimensional folded site frequency spectrum (SFS) was used to calculate Fst using the Bhatia estimator (ANGSD v. 0.940, realSFS fst index; realSFS fst stats2; Korneliussen et al., 2014). To calculate dxy, we first re-calculated the 2D SFS for every 50kb window, then calculated dxy using a modified version of dxy_wsfs.py script from (D. Marques, https://github.com/marqueda/PopGenCode/blob/master/dxy_wsfs.py, accessed Nov. 2023). The script was modified to run in R (dxy_wsfs.R; supplementary materials). We also used ANGSD to calculate π and Tajima’s D over 50kb windows, and calculated the difference in nucleotide diversity (Δπ) and Tajima’s D (ΔD) as πCH - πCO and DCH-DCO respectively. Outlier windows for each metric were identified as windows more than 3x the standard deviation above (Fst, Dxy, Δπ, ΔD) or below (Δπ, ΔD) the mean.

We intersected gene loci (+/-2kb, described above) with 50kb windows (+/-2kb, described above) and used hypergeometric tests in R (stats::phyper ()) to evaluate whether differentially expressed genes were more likely to appear in regions of high differentiation or divergence than expected by chance. We further tested whether the relationship between expression and genomic divergence differed inside and outside of the putative inversion using Kruskall-Wallis and Dunn’s multiple comparison tests (rstatix, v.0.7.2; Kassambara, 2021). Although we did not calculate population genomic metrics for individual genes, we nevertheless ran a second set of tests as a control, using a subset of non-differentially expressed genes matched for length with the differentially expressed genes (nullranges, v.1.2.0; M. Love et al., 2022), as gene length covaries with some population genomic metrics.

## Results

Ecotype was a major driver of gene expression differences in *T. conura* larvae. The CH and CO ecotypes separated on the first and second axes in a principal component (PC) analysis of the 500 genes with the most variable expression (Figure 1F). In contrast, larvae did not cluster by cross-fostering treatment on any of the first 5 PC axes (Figure S4). Consistently, hierarchical clustering using all genes grouped larvae by ecotype, especially CO larvae, with some evidence that cross-fostered larvae tended to cluster together with the CO ecotype, regardless of whether they were cross fostered to the natal or novel host (Figure S5). Jointly, these expression patterns suggest that evolved expression differences affect a much larger portion of the transcriptome than plastic responses to the cross-fostering treatment.

### Differential expression between ecotypes

Larvae of the CH and CO ecotypes use different genes to deal with toxic or harmful components in their host plants. Comparing the control lines H and O, we detected 488 genes upregulated in H larvae and 445 genes upregulated in O larvae, which together make up 7.97% of the filtered, expressed transcriptome (Figure 2A). Both gene sets were primarily enriched for GO terms involved in regulation of biological processes and metabolism. Detoxification (GO:0098754, Fisher’s exact test with parent-child algorithm p = 7.6×10^−5^), cellular detoxification (GO:1990748, p = 3.1×10^−5^), and cellular response to toxic substance (GO:0097237, p = 3.5×10^−3^), were all explicitly enriched in the genes upregulated in O larvae. This gene set was also enriched for biosynthesis and metabolism of glycoproteins (Table S3), which are involved in herbivorous insect responses to plant chemical defenses like flavonoids (Aurade et al., 2011; Dermauw & Van Leeuwen, 2014), which are the dominant chemical defense in thistles (Jordon-Thaden & Louda, 2003). Genes upregulated in control line O larvae were also enriched for protein folding (GO:0006457, p = 9.1×10^−5^) and the endoplasmic reticulum unfolded protein response (GO:0034975, p = 6.0×10^−3^), which are both characteristic of the insect integrated stress response (Rosche et al., 2021). Genes upregulated in H larvae were highly enriched for various metabolic processes (e.g., GO:0080090, p = 3.2×10^−11^), and marginally enriched for toxic responses, such as toxin transport (GO:1901998, p = 3.3.x10-2), and response to chemical stimulus (GO:0070887; p = 2.3×10^−3^) and chemical stress (GO:0062197; p =4.7×10^−2^). Together, these differentially expressed genes confirm that C. oleraceum and C. heterophyllum represent distinctly different chemical and nutritional environments that CH and CO ecotype larvae have evolved different molecular mechanisms enabling them to feed on these host plants.

**Figure 2.**
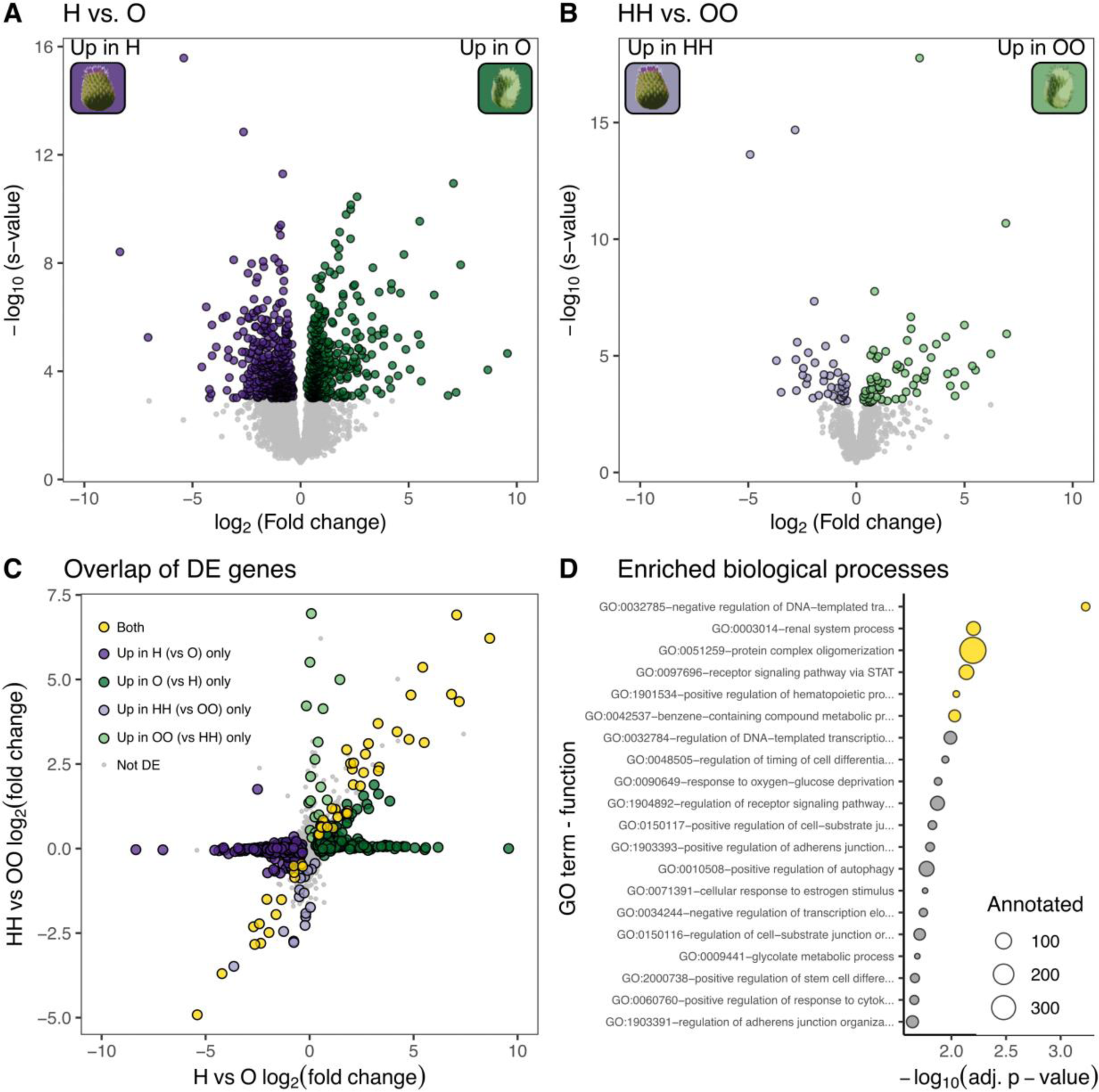
Differential expression between CH and CO larvae. Differentially expressed genes (A) between control lines (H vs. O) and (B) between larvae cross-fostered to their natal host (HH vs. OO). Differential expression was tested using Wald’s Chi squared tests and significantly differentially expressed genes had an s-value < 0.001 after applying a log_2_ fold change shrinkage estimator. (C) Overlap between differentially expressed gene sets in A and B. Genes that are significantly differentially expressed in both comparisons are shown in yellow. (D) Biological process enriched (p < 0.01, yellow) in the subset of overlapping genes in A and B. Enrichment was calculated using Fisher’s exact test adjusted with the parent-child algorithm and p-values were corrected for multiple testing using an FDR correction.

In comparison, many fewer genes were differentially expressed between larvae of the two ecotypes when cross-fostered to their natal hosts (HH vs. OO; Figure 2B). This is largely due to high variation within each of the sample groups, likely introduced by the stress of the cross fostering process. Overall, 40 genes were upregulated in HH larvae and 64 genes were upregulated in OO larvae, and only 53 of these genes overlapped with the 933 genes that were differentially expressed between control larvae (Figure 2C). These 53 genes that differed between host races on their native plants regardless of cross-fostering were enriched for a small set of functions that reveal the consistent differences between CH and CO specialist larvae feeding on their natal hosts (Figure 2D, Table S4), including renal system processes (GO:0003014, rank = 2, p-value = 6.3×10^−3^), which in dipterans implicates structures like nephrocytes and malpighian tubules that are involved in fluid balance, electrolyte balance, pH balance, and disposal of toxic waste products (Denholm & Skaer, 2009; J. Xu et al., 2022). Importantly, these shared genes were also enriched for metabolism of benzene-containing compounds (GO:0042537, rank = 6, p-value = 6.0×10^−3^). Flavonoids are composed of two benzene rings linked by a three-carbon pyran ring (Celeste-Dias et al. 2021). This result further supports that the divergent defensive flavonoid chemical profiles of *C. heterophyllum* and *C. oleraceum* play a role in different expression and performance of specialist larvae on these host plants.

### Expression plasticity in ancestral and derived ecotypes

We tested to which extent larvae can plastically shift gene expression in response to a different host, and whether this ability differs between the ancestral (CH) and derived (CO) ecotypes. Overall, larvae that were cross-fostered to a novel host had very few differentially expressed genes compared to larvae cross-fostered to their natal host, with fewer than 10 genes differentially expressed in ancestral (HH vs. HO) and derived (OO vs. OH) larvae (Figure S6A, B). Nevertheless, even a small number of differentially expressed genes could represent adaptive transcriptional plasticity. We predicted that genes with adaptive plastic expression in larvae cross-fostered to novel hosts would match the expression differences found between the two ecotypes. For example, if gene expression is adaptively plastic, differences between HH and HO larvae should match those between HH and OO larvae. However, we found only a single gene matching this pattern for each of the ecotypes (Figure S6 C,D). In the ancestral CH ecotype, this gene was Tcon_g9682, aka RARS, which belongs to the class-I aminoacyl-tRNA synthetase family. Expression of RARS was elevated in HO larvae and all CO ecotype larvae (Figure S6C inset). In the derived CO ecotype, Tcon_g14248 (TRAPCC5, a trafficking protein particle complex) matched the expected pattern of expression based on host plant rather than ecotype. However, when we further investigated TRAPPC5 expression across treatments, we found that this gene was not in the subset of shared DE genes between ecotypes (Figure 2C). Rather, control H larvae and cross-fostered HH larvae had very different expression levels of TRAPPC5, as did O and OO larvae (Figure S6D inset). These results could suggest that neither ecotype has the capacity for extensive, adaptive transcriptional plasticity when switched to a novel host during the third instar.

### Ecotype-, stress- and plasticity-associated patterns in coexpressed genes

As an alternative test of evolved and plastic differences in gene expression across treatments, we used weighted gene coexpression network analysis to identify 21 modules of coexpressed genes (Figure S7A, Table S5). We further tested for modules that were meaningfully correlated (r > 0.5) and significantly associated (adj. p < 0.01) with predicted expression patterns (Figure S3). This was done by creating binary vectors for the samples reflecting each of five possible scenarios, including ecotype, cross-fostering, reciprocal plasticity, and asymmetric plasticity in either the CH or CO ecotype (Figure S3). These vectors were then correlated with eigengene expression, which is a summary of expression for all genes in that module in that sample, for each of the modules. Thus a significantly positively correlated module is a module in which eigengene expression was consistently high for samples assigned 1 in a given scenario, and low for samples assigned 0.

We found that one third of the modules were correlated with larval ecotype (Figure S7B, Figure 3A). None of the modules were expressed as expected under reciprocal plasticity, where larval expression reflects the host plant they were feeding on regardless of ecotype. Similarly, no modules were expressed in a manner consistent with CO plasticity, where CO larvae cross-fostered to *C. heterophyllum* have CH-like expression. However, three modules had a significant signature of CH plasticity, with CH larvae cross-fostered to *C. oleraceum* having CO-like expression.

**Figure 3.**
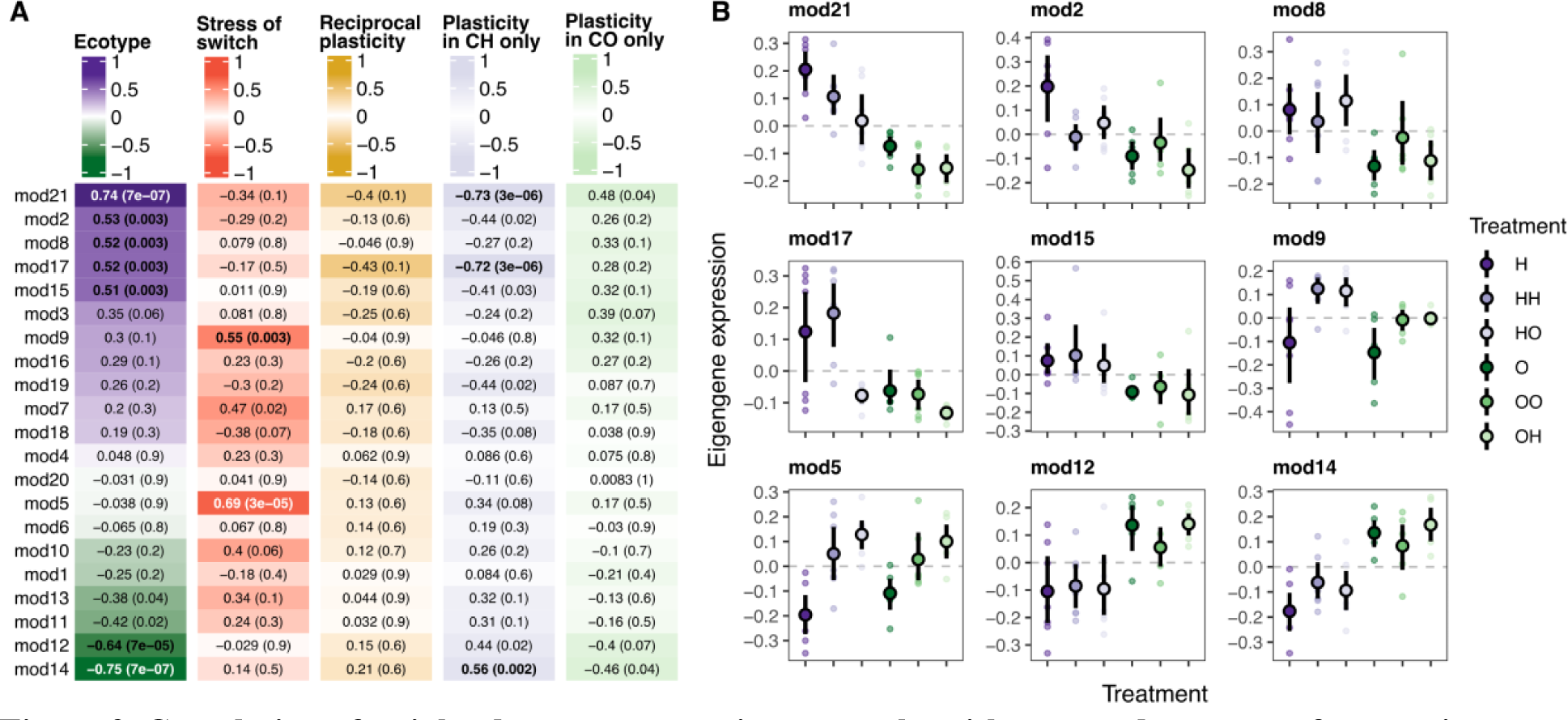
Correlation of weighted gene coexpression networks with expected patterns of expression across treatments. (A) Heatmaps showing correlation of module eigengenes with ecotype (1 = CH, 0 = CO), stress of cross-fostering/switch (1 = switched, 0 = not switched), reciprocal plasticity (1 = feeding on *C. oleraceum*, 0 = feeding on *C. heterophyllum*), plasticity in CH only (1 = CO ecotype or HO treatment, 0 = CH feeding on *C. heterophyllum*), plasticity in CO only (1 = CH ecotype or OH treatment, 0 = CO feeding on *C. oleraceum*). Pearson’s correlation coefficient and the adjusted student asymptotic p-value are in bold if they meet our threshold for a meaningful, significant correlation (r > 0.5 & p < 0.01; see Figure S3). (B) Expression of eigengenes (first principal component of gene expression) across treatments for modules with significant correlations in A. Small points show eigengene expression for each sample, and black points and vertical lines represent means and 95% confidence intervals, respectively.

Of the modules associated with CH plasticity, module 17 (n = 48 genes) was most strongly correlated (r = −0.72, p = 3×10^−6^). Genes in module 17 had low expression in any larva feeding on *C. oleraceum*, and highly variable expression in H and HH larvae (Figure 3B, S7). This module was overwhelmingly enriched for reproductive processes (Figure S8D, Figure S9D), suggesting CO larvae may have a delayed life history progression relative to CH larvae of the same size, and that feeding on *C. oleraceum* either directly or indirectly (e.g., by slowing growth) delays the onset of reproductive development in larvae of the CH ecotype.

Module 21 (n = 38 genes) expression was also significantly correlated with CH plasticity (r = - 0.73, p = 3×10^−6^), but was also the module most correlated with the CH ecotype overall (r = 0.74, p = 7×10^−7^). Overall, genes in this module were less expressed in the CO ecotype, but cross-fostered larvae from the CH ecotype also tended to express lower levels of these genes. This module was enriched for cell cycle processes, such centriole assembly (GO:0098534, rank = 1, p = 5.2×10^−4^) and replication (GO:0007099, rank = 2, p = 6.2×10^−4^), chromatin (GO:0006325, rank = 3, p = 1.4×10^−3^) and microtubule organizing center (GO:0031023; rank = 6, p = 2.7×10^−3^) organization (Figure S8A, Figure S9A). Further, this module was enriched for salivary gland histolysis (GO:0035070, rank = 8, p = 5.0×10^−3^), a specific process of salivary gland breakdown during dipteran metamorphosis (de Cassia Santos Przepiura et al., 2020). Together with module 17, this supports a faster life history in CH flies that is significantly slowed by cross-fostering in *C. oleraceum* buds.

Finally, module 14 (n = 618 genes) was the third module correlated with CH plasticity (r = 0.56, p = 0.002; Figure 3A). However, assessed visually, eigengenes for this module do not fully match our expectations for CH plasticity (i.e., Figure S3), and overall, the module was also highly correlated with ecotype (r = −0.75, p = 7×10^−7^), which likely influenced the correlation with CH plasticity. However, this module contains RARS (Tcon_9682) which we previously identified as plastically expressed in HO flies using our differential expression approach. The module was also enriched for numerous processes involved in cellular respiration, detoxification and protein transport (Figure S8, Figures S9).

Two modules were significantly correlated with a stress effect from cross-fostering:, module 5 (n = 35 genes, r = 0.69, p = 3×10^−5^) and module 9 (n = 45 genes, r = 0.55, p =0.003). In these modules, expression in the cross-fostered larvae was consistently higher than control larvae. This stress effect was corroborated by significant functional enrichment in module 5 of regulation of metabolic and catabolic processes (GO:0019222, rank = 3, p = 2.9×10^−3^; GO0031323, rank = 7, p = 5.3×10^−3^; GO:0031329, rank = 1, p = 2.2×10^−3^; GO:0009894, rank = 4, p = 3.8×10^−3^; Figure S8F) and cytokine-like and interferon signaling and response (GO:0060338, rank = 5, p = 4.2×10^−3;^ GO:0060759, rank = 30, p = 2.2×10^−2^), which are associated with immune system responses (Labropoulou et al., 2024; Rosche et al., 2021). Module 9 was enriched for midgut development (GO:0055123, GO:0007496, GO:0007494) and endothelial cell development and regeneration (Figure S8G). The functional enrichments of these modules suggest that the process of moving to a new host is metabolically and immunologically stressful, regardless of whether the new host is the natal, adapted host or the novel, challenging host.

### Expression differences associated with genomic architecture and population divergence

To uncover the relationship between expression differences and genomic architecture we assess distribution of differentially expressed genes in relation to genetic divergence. Nilsson et al. (2023) identified a large putative inversion (∼104Mb, 83 contigs) on LG3 in the *T. conura* assembly as the major locus of genomic divergence between the ecotypes in parallel contact zones east and west of the Baltic Sea. Here, we focus only on the two source populations for our cross-fostered larvae (Figure 1B) to test whether constitutive or plastic expression differences tended to be associated with highly differentiated or divergent regions inside and outside of this inversion. As expected, we identify a large peak in Fst and dxy at the putative inversion (Figure 4A,B). This region is also characterized by a positive peak in Δπ (nucleotide diversity higher in the CH ecotype than the CO ecotype, Figure S10) and a negative ΔD (Tajima’s D lower in the CH ecotype than the CO ecotype, Figure S10; Figure 4C, D). Using these two populations, we identified a subset of genomic windows that were consistently supported by differentiation, divergence and selection metrics (> mean ± 3 std. deviations in 3 out of 4 metrics; Figure 4F). These windows contained 129 genes (Figure 4F) and were overwhelmingly located within the putative inversion. Only one gene, Tcon_g8259 (unnamed in our functional annotation, possible ortholog of Dmel\CG13928 BLAST score = 63.1586 bits), was located in outlier windows outside of the inversion.

**Figure 4.**
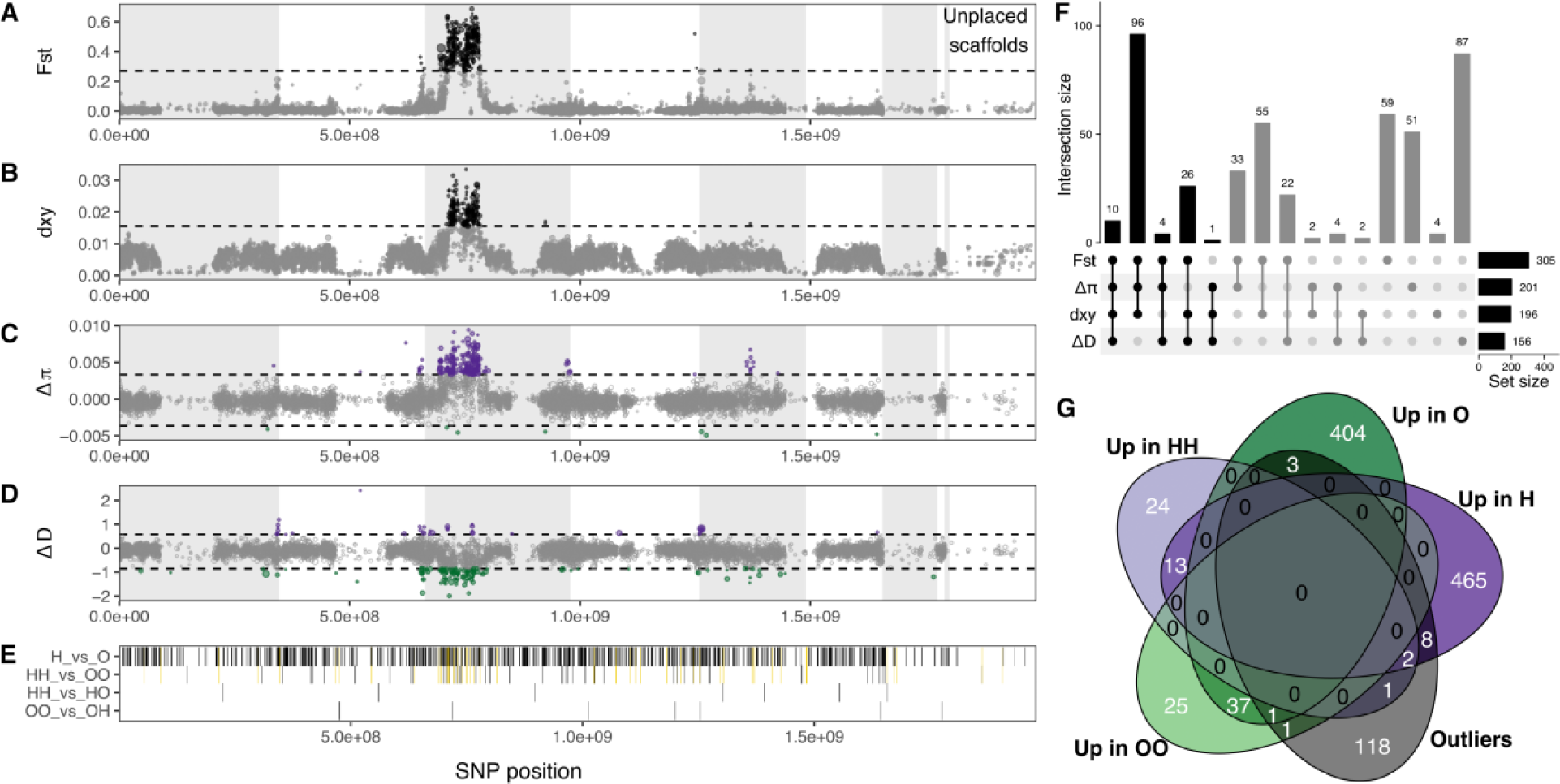
Overlap between differentially expressed genes and regions of high genomic differentiation and divergence between CH and CO populations. (A) Fst (Bhatia estimator) and (B) dxy calculated from the allele frequency spectrum over 50kb windows based on the CO and CH populations used for the cross-fostering experiments (Figure 1D). Horizontal dashed lines represent the mean + 3 standard deviations. Outliers are colored in black. (C) Nucleotide diversity difference (Δπ) and (D) Tajima’s D difference (ΔD) between CH and CO populations. Horizontal dashed lines represent the mean +/- 3 standard deviations, and outliers are colored in purple if π or D were higher in CH, green if higher in CO. Windows with lower than 20% coverage were excluded, resulting in gaps along the ordered contigs. Contigs are ordered according to hypothetical linkage groups. Linkage groups are delineated as light gray or white bands, with the rightmost white band containing unscaffolded, low coverage contigs. (E) Locations of genes that were differentially expressed between treatments were distributed throughout the genome. Yellow bands show genes that overlap between H vs. O and HH vs. OO (Figure 2C). (F) UpsetR plot of genes falling in outlier windows for each metric. We classified outlier genes as those overlapping outlier windows in 3 out of 4 population genomic metrics (black bars). (G) Venn diagram visualizing overlap between differentially expressed genes (H vs. O and HH vs OO) and outlier genes.

We next tested whether genes that were differentially expressed between ecotypes were also likely to overlap these regions of high differentiation and divergence. Both the set of genes that were differentially expressed between HH and OO, and the subset of these that were also differentially expressed between H and O (i.e., “both”, Figure 2C, yellow lines Figure 4E), overlapped with outlier genes (i.e., Figure 4F) significantly more than expected by chance (HH vs OO odds ratio = 4.49, hypergeometric test, adj. p-value = 0.011; “both” odds ratio = 5.27, hypergeometric test, adj. p-value = 0.028). However, when we compared gene sets from H and O to the set of outlier genes (Figure 4F), differentially expressed genes overlapped no more than expected by chance (Figure 4G, odds ratio = 1.35, hypergeometric test, adj. p-value > 0.999). None of the genes that were differentially expressed between HH and HO or between OO and OH overlapped with outlier windows (Figure 4E).

We investigated if there was evidence for increased differentiation, divergence, or selection for differentially expressed genes by assessing if Fst, dxy, Δπ, and ΔD, π or Tajima’s D differed between genes that were or were not differentially expressed. This analysis was performed separately for genes inside and outside of the inversion. Overall, genes within the inversion that were differentially expressed between ecotypes (H vs. O) were not significantly different from genes that were not differentially expressed, apart from marginally significantly lower nucleotide diversity in the CO population (Figure S11; Table S7). However, among the genes outside of the inversion, differentially expressed genes were less divergent (lower dxy; Figure S11B) and accordingly had significantly lower π in both populations (Figure S11E,F). This pattern was maintained when we subset the non-differentially expressed genes to match the gene length distribution of the differentially expressed genes, and may result from purifying selection on or near genes that are functionally important for host plant use. No significant differences were found when we focused on genes that were differentially expressed in HH vs OO (Figure S12; Table S8).

Finally, we also compared these metrics among the expression modules, excluding module 20 which only contained 7 genes after membership filtering (Figure S13). While there were some overall patterns (e.g., dxy and π in both populations tended to be higher in modules 1, 8, 10 and 19, and lower in modules 5, 13,14, and 16, Dunn’s multiple comparison, Table S9), these patterns did not align with module correlations with ecotype, stress or plasticity (Figure 3A).

## Discussion

To what extent evolution of gene expression enables adaptation to novel environments, and how altered gene expression interacts with genetic divergence and altered genomic architecture during this process is crucial for understanding the adaptive potential of organisms (Ballinger et al., 2023; López-Maury et al., 2008; Pavey et al., 2010; Triant et al., 2021). Importantly, whether the changes in gene expression that enable the use of novel environments are evolved or plastic has consequences for the potential to retain the ability to use ancestral environments (Celorio-Mancera et al., 2023; Ho et al., 2020; Ho & Zhang, 2018). Moreover, few studies have addressed whether the process of adaptation to a novel niche affects the capacity for transcriptional plasticity and subsequent broad niche use.

Here we investigate evolved and plastic changes in gene expression underlying ecotype adaptation to a novel host plant in the peacock fly, *T. conura*. Overall, we found limited evidence for adaptive transcriptional plasticity in response to feeding on the alternate host in either the ancestral CH ecotype and derived CO ecotype. Nevertheless, coexpression analysis uncovered three modules of genes for which expression in CH larvae feeding on *C. oleraceum* shifted to match expression of the CO ecotype, providing some evidence for the hypothesis that transcriptional plasticity would be greater in the ancestral than the derived ecotype. In contrast, we found extensive constitutive transcriptional differences between the two ecotypes, many of which were robust to the stressful cross-fostering treatment. Genes that were differentially expressed in the two ecotypes were enriched for metabolism and detoxification functions, likely reflecting adaptations to differences in the nutritional and chemical profiles of *C. heterophyllum* and *C. oleraceum.* Finally, we found that these constitutively differentially expressed genes were more likely than chance to to fall in highly divergent regions of the genome, specifically within the large putative inversion on LG3. Together these results suggest that gene expression evolution has contributed significantly to adaptive phenotypic divergence between CH and CO lineages and genomic architecture is playing an important role in shaping the transcriptomes of these ecotypes.

### Gene expression divergence between ecotypes

Host plant chemistry is a major driver of host plant specialization and diversification in insects (Birnbaum & Abbot, 2020; Jaenike, 1990; Jousselin & Elias, 2019; van der Linden et al., 2021). Our results support a critical role for thistle chemistry in the adaptive divergence of gene expression between the CH and CO ecotypes of *T. conura*. Broadly, we found that differentially expressed genes between the ecotypes are significantly enriched for biological processes related to detoxification and responses to toxic chemicals, suggesting that the ecotypes use highly divergent molecular mechanisms to confront the challenges of feeding on their respective preferred hosts. Specifically, genes that were differentially expressed both between the control H and O treatments and between HH and OO were especially enriched for metabolism of benzene-containing compounds. The dominant class of secondary metabolites produced by thistles are flavonoids (Jordan-Thaden & Louda 2003), which are composed of two benzene rings linked by a three-carbon pyran ring (Dias *et al*., 2021). Flavonoids have been shown to have either deterrent or even toxic effects on herbivorous insects or oviposition attractant or feeding stimulant effects (Aurade *et al*., 2011; Mierziak *et al*., 2014; Dias *et al*., 2021). Functional enrichment of metabolism of benzene-containing compounds suggests the flavonoid chemical profiles of *C. heterophyllum* and *C. oleraceum* are important selection pressures acting on gene expression in specialist larvae on these host plants.

Evolved differences in gene expression between the CH and CO ecotypes likely act as a barrier to gene flow between the ecotypes. Larval feeding assays have shown that both ecotypes have significantly reducedsurvival on the wrong host plant (Diegisser et al., 2008). Our results suggest that evolved differences in gene expression, adapted to the different chemical and possibly nutritional profiles of the host plants, are the molecular mechanisms underlying these fitness costs. Limited plasticity, especially in the derived CO ecotype, means that larvae are not buffered against maladaptive oviposition decisions of adult females, and there should be strong selection against laying eggs in the wrong host plants in parapatric and sympatric contact zones. Because mating takes place on the host plant (Romstöck & Arnold, 1987), increased selection on host recognition traits is likely to limit not only maladaptive oviposition but also encounters between males and females of the different ecotypes.

We expect divergence in both *cis-* and *trans-* regulatory elements to contribute to the extensive expression differences uncovered between the ecotypes. For example, many of the genes found to be both differentially expressed and highly genetically divergent between ecotypes were annotated with regulatory roles, including transcription factors and methyltransferases involved in epigenetic modifications for active gene transcription (Table S6). Divergence in evolved regulatory networks has the potential to lead to transgressive phenotypes in hybrids that, while not intrinsically inviable in the classic sense of Batesian-Dobzhansky-Muller incompatibilities, may in turn lcause reduced fitness in available niches (Thompson et al., 2023). While we have yet to explore gene expression and fitness in F1 hybrids or subsequent backcrosses, hybridization could produce combinations of *cis-* and *trans-* regulatory elements that disrupt the host-specific gene expression profiles we have uncovered in this paper, leading to reduced hybrid fitness.

### Genomic architecture of differential gene expression

Our results suggest a significant role for the large inversion on LG3 identified by Nilsson et al. (2023) in gene expression divergence, as the inverted region was significantly enriched for genes that were consistently differentially expressed between the ecotypes. Inversions are well known to be important drivers of ecological divergence and speciation (Berdan et al., 2023; Faria et al., 2019; Feder & Nosil, 2009; Fuller et al., 2016; Westram et al., 2022) and have been identified in a number of systems exhibiting rapid local adaptation to and divergence in novel niches (Ayala et al., 2013; Koch et al., 2021; Kollar et al., 2023; Lee et al., 2017; Lowry & Willis, 2010; Morales et al., 2019; Twyford & Friedman, 2015). In particular, inversions have been found to underlie ecological divergence and reproductive isolation in several well known examples of host-plant associated differentiation and host race formation. Inversions segregate between apple- and hawthorn-infesting populations of another Tephritid fly, *Rhagoletis pomonella* (Egan et al., 2015; Feder & Nosil, 2009; Ragland et al., 2017). Large inversions also underlie both a host-associated color polymorphism (Nosil et al., 2018) and inter- and intra-specific variation in feeding on redwood trees by *Timema* walking sticks(Nosil et al., 2023). However, experimental evidence for how inversions alter gene expression during local adaptation and divergence is still limited (Berdan et al., 2023).

Inversions can influence gene expression in two main ways (Berdan et al., 2023). First, inversion breakpoints can break genes or rearrange the structural relationships between regulatory elements and the protein coding regions of genes. Even though this is only likely to impact a small fraction of genes, depending on their roles within larger gene regulatory networks, genes at or near breakpoints could have cascading effects on gene expression throughout the genome.

Second, as regions of low recombination, inversions can capture and accumulate divergent sequence variation more quickly than collinear regions of the genome (Faria et al., 2019; Schaal et al., 2022), resulting in rapid evolution of both *cis-*regulatory elements and coding regions of *trans-*acting factors located within the inversion. For example, 80.6% of genes that were differentially expressed between inversion genotypes of the seaweed fly *Coelopa frigida* were located within the inversion (Berdan et al., 2021), suggesting a major role for *cis-*regulatory sequence divergence within the inversion. Which of these mechanisms are more likely to be happening in the *T. conura* system is still unclear. We are currently unable to assess breakpoint dynamics due to lack of contiguity in our *T. conura* assembly. However, we have extensive evidence for sequence divergence between the inversion haplotypes resulting from reduced introgression and positive selection, especially on the CO ecotype (Nilsson et al. 2023). Moreover, windows containing differentially expressed genes tended to have lower nucleotide diversity in both populations regardless of whether they were found inside the inversion or not. Nevertheless, we did not find differences Fst, dxy, or Tajima’s D in windows around differentially expressed genes compared to those that were not differentially expressed. Thus, while we hypothesize that sequence divergence within the inversion is contributing to the large number of differentially expressed genes between the ecotypes, we were unable to identify specific signatures of divergence or selection supporting this hypothesis.

### Limited, asymmetric expression plasticity

Based on the oscillation and flexible stem hypotheses, we hypothesized that transcriptional plasticity in response to novel host plants should be common (Celorio-Mancera et al., 2023; Gibert, 2017), but this was not the case. We also expected that if the CO ecotype experienced strong selection on gene expression when colonizing and adapting to *C. oleraceum*, this should have led to genetic assimilation, resulting in asymmetrically higher plasticity in the ancestral CH ecotype. While overall evidence for plasticity was limited, our results broadly fit these predictions: three gene coexpression modules exhibited patterns of gene expression plasticity unique to CH larvae, while no modules exhibited patterns consistent with plasticity unique to CO larvae. However, what remains unclear is whether the asymmetric plasticity observed in modules 21, 17 and 14 is adaptive, neutral or even nonadaptive. In order for plasticity to be adaptive, it should contribute to increased fitness in the new environment, and expression should match, or approach, the optimum for that environment (Ghalambor et al., 2015). And yet, expression matching may not always be adaptive. For example, module 17 genes, which were found to be enriched for reproductive development and maturation functions, were only expressed in noticeable amounts in H and HH larvae, and were negligibly expressed in all CO larvae as well as CH larvae feeding on *C. oleraceum* (HO). All larvae in our study were size-matched, but it is possible that reproductive development starts at a smaller size in CH than CO larvae, and rather than matching an adaptive expression optimum in CO larvae, larvae in the HO treatment experienced delayed or stunted development by feeding on *C. oleraceum*, even for a short period of time. Under this scenario, the “CH-plasticity” of module 17 genes may represent nonadaptive changes in gene expression that match CO larval expression by chance. Similarly, module 21 genes were enriched for developmental functions, such as cell-cycle regulation and salivary histolysis, both of which could be associated with pre-pupal development, and could have been inhibited by feeding on a suboptimal host plant.

We did not find any evidence for reciprocal plasticity, in which both ecotypes share host-plant specific transcriptional patterns. One prediction of the oscillation hypothesis is that shared molecular mechanisms facilitate colonization and recolonization of novel and ancestral host plants (Celorio-Mancera et al., 2023; Ho et al., 2020). Accordingly, Celorio-Mancera et al. (2023)found that coexpressed gene modules were shared between generalist and specialist species of *Polygonia* and *Nymphailis* butterflies and similarly expressed when feeding on the same host plant. In contrast to these findings, we do not find evidence for gene expression activated depending on host plant use in the CH and CO *T. conura* ecotypes.

In contrast to examples in generalist butterflies, the focal *T. conura* populations are highly specialized. However, both ecotypes currently have populations using multiple host plants within their distribution. CH flies oviposit and successfully develop in both *C. heterophyllum* and *C. palustre* in the northern British Isles (Diegisser et al., 2009; Romstöck-Völkl, 1997), and CO flies oviposit in *C. oleraceum* and *C. acaule*, among others, in several regions of the Alps (Romstock-Vokl 1997). Transcriptional plasticity is known to be important for single populations using multiple hosts (Birnbaum & Abbot, 2020), depending on the chemical similarity of the plants (Celorio-Mancera et al., 2016). Whether these oligophagous populations would show greater transcriptional plasticity when cross-fostered on *C. heterophyllum* and *C. oleraceum* is not known, and could be a useful next step for understanding the roles of adaptive plastic phenotypes and their role in colonizing new hosts.

The importance of transcriptional plasticity in host use is a difficult question to address because larval age, experience and length of exposure can all interact to affect larval gene expression profiles (Birnbaum & Abbot, 2020; Schneider et al., In prep). One potential concern of our study design is that larvae were only allowed to feed on the novel host plant for 6 hours, which may not be long enough to detect a consistent, regulated change in gene expression on the new host. For example, Schneider et al. (In prep) found that after two hours of feeding on a new host, gene expression in larvae of the butterfly *Polygonia c-album* was still best explained by the natal host. It was only after 17 hours that a main effect of the second host plant on gene expression could be detected. Despite this concern, our results clearly show that *T. conura* larvae of both ecotypes are able to rapidly alter gene expression in the 6 hour time frame. We detected two small coexpressed modules that were consistently upregulated in response to the acute stress of the cross-fostering design. These modules were strongly enriched for functions involved in the insect integrated stress response (Harding et al., 2003; Rosche et al., 2021). It is also unclear how transcriptional plasticity changes over the lifespan of a developing insect, and how this transcriptional plasticity translates to phenotypic plasticity and fitness. Potentially, larvae exhibit plasticity early in development which is gradually lost over longer exposure to the same host environment. To our knowledge, there have been no studies on how host-associated transcriptional plasticity changes through juvenile and adult development, (but see Celorio-Mancera et al., 2013). However, phenotypic assays suggest that holometabolous larvae often habituate to their natal feeding environment, altering subsequent acceptance (Huang & Renwick, 1995) or performance (Söderlind et al., 2012). How these changes in phenotypic plasticity across life stages could relate to underlying transcriptional plasticity is an interesting topic for future research.

### Conclusions and future perspectives

Overall, this study contributes to our understanding of the role of ancestral plasticity and genetic architecture in evolved changes in gene expression enabling adaptation to novel environments. While there is ample evidence that plastic changes to gene expression are important for the use of multiple host plants (Birnbaum & Abbot, 2020), the extent to which these changes translate into evolved differences in gene expression between specialists remains unclear. We find limited evidence for plasticity, and while the ancestral ecotype altered more genes in response to exposure to the alternate host plant, these genes and modules were not clearly related to the selective environment of the derived host. Thus it is unclear whether these changes represented adaptive plasticity in the ancestral ecotype. Instead, genes involved in processing host specific chemicals are constitutively differentially expressed between ecotypes. Interestingly, our findings support the role of genomic architecture in the ecotype-specific gene expression profiles, as consistently differentially expressed genes were more densely located within the large, putative inversion on LG3. Whether the overrepresentation of differentially expressed genes within the inversion is due break-point induced dynamics or result from selection acting within the inversion is an outstanding question for future studies. Here, we provide an initial look at the role of transcriptional divergence and plasticity in ecotype formation and incipient speciation of *T. conura.* The *T. conura* genome is highly repeat rich, with a large expansion of DNA and retrotransposons (Nilsson et al., 2023). Examining the roles of transposable elements in addition to small and non-coding RNAs and alternative splicing in molding the ecotype specific patterns of gene expression and plasticity is likely to further improve our understanding of the mechanistic basis enabling the use of a novel host.

## Acknowledgements

We thank Kalle Nilsson and Emma Kärrnäs for assistance in the field and Oliver Moss for training in RNA extraction. This work was funded by a Wenner-Gren fellowship, a Swedish research council starting grant, as well as funding from the Crafoord foundation, Erik Philip Sörensens stiftelse and Carl Tryggers stiftelse to A.R. and from the Royal Physiographic Society in Lund to J.O. The authors acknowledge support from the National Genomics Infrastructure in Genomics Production Stockholm funded by Science for Life Laboratory, the Knut and Alice Wallenberg Foundation and the Swedish Research Council, and SNIC/Uppsala Multidisciplinary Center for Advanced Computational Science for assistance with massively parallel sequencing and access to the UPPMAX computational infrastructure.

## Data availability

Upon publication, RNA sequence data will be made available through the European Nucleotide Archive (PRJEB73529). Gene predictions and functional annotations, as well as all scripts necessary for analysis and visualization of the data will be archived and made public on GitHub.

