## Supplementary Materials for "Evolved and plastic gene expression in adaptation to a novel niche"

### Supplementary Methods

#### *Cross-fostering*

Cross-fostered larvae were moved into uninfested buds (<20 mm) of either the natal or novel host. The bud was first cut in half, leaving the base of each half attached at the peduncle so that the bud remained connected to the plant. A small cavity (~4mm) was created in the uninfested bud to give space to the cross-fostered larva. After placement of the larvae, the halves were placed back together and wrapped in parafilm to minimize moisture loss. It is possible that damaging the thistle buds in this way affected the induced secondary metabolites in the thistle tissue. However, there is currently very little information about the constitutive or induced defensive chemicals in these plants (Jordon-Thaden & Louda, 2003). Larvae were placed into new buds in a split brood design, with one individual from each clutch assigned to each treatment. Difficulties with larval survival, RNA extraction and sequencing introduced some variation into the split brood design, and overall, we sampled 6-7 larvae in each treatment from a total of 11 families (Table S2).

#### *Gene prediction and annotation with BRAKER3*

Trimmed reads were used to generate a new, improved gene annotation for the *T. conura* genome. Reads were mapped to the genome using the splice-aware aligner HISAT2 (REF) with the –dta flag activated. BRAKER3 (Gabriel et al., 2023) was used to generate a gene annotation, guided by proteins from Arthropoda ODB_11 database (Kuznetsov et al., 2023) and the aligned RNAseq reads, excluding samples with poor mapping rates. We also included RNAseq reads from pupal and adult stages of both ecotypes for a more complete set of gene predictions. We used the default BRAKER3 pipeline, but a large proportion of single exon genes motivated us to re-merge the protein-based and the RNA-based annotations with TSEBRA (Gabriel et al., 2021), reducing the support necessary for introns from 1 to 0.25. The result was a highly complete annotation (BUSCO v. 5.3.1; Manni et al., 2021): 98.0% complete (S: 73.6%, D: 24.4%), 0.8% fragmented, 1.2% missing, diptera_ODB10, n = 3285, Supplementary Figure S1), with a total of 25175 genes and 30632 transcripts.

The coding sequences from this version of the annotation were used to quantify expression of *T. conura* transcripts (see below). Coding sequences were also converted to amino acid sequences with gffread (v. 0.12.7; Pertea & Pertea, 2020) and submitted to EggNOG mapper (v. 2.1.12; http://eggnog-mapper.embl.de/; Cantalapiedra et al., 2021; Huerta-Cepas et al., 2019) for functional annotation. We produced two functional annotations, one in which taxonomic scope was set to 'default', which adjusts scope based on each protein sequence, and one in which taxonomic scope was set to 'arthropoda'. The first is better for gene set enrichment analysis because overall more genes are assigned GO terms. The second is better for assigning gene names and functions, as the terminology is comparable to other arthropod taxa (e.g., *Drosophila melanogaster*).

We assigned gene IDs following TSEBRA's default scheme, with the species prefix "Tcon" appended to a gene number ("g1"), followed by a transcript number ("t1"), for example "Tcon_g1.t1"). We refer to genes using this nomenclature throughout the manuscript, and use the gene name assigned by the EggNOG functional annotation when available.

To capture the extent of coding and noncoding regions, we added untranslated regions (UTRs) using the BRAKER3 script stringtie2utr.py, per author recommendations (K. Hoff; https://github.com/Gaius-Augustus/BRAKER/issues/638#issuecomment-1741029025). We additionally converted the GTF with UTRs into a BED file (AGAT, agat_convert_sp_gtf2bed.pl; Dainat et al., 2022) and added a 2kb window around each locus to capture promoter and other regulatory regions (BEDTools slop v.; Quinlan Laboratory, 2023). This BED file was intersected (BEDTools Intersect) with 50kb windows over which we estimated differentiation, divergence, nucleotide diversity and Tajima's D for the CH and CO populations (see below).

### Supplementary Tables

***Table S1. Treatment design and sample sizes for* Tephritis conura *ecotypes feeding on* Cirsium *thistle host plants.***

Treatment codes correspond to design outlined in Figure 1. One sample was excluded based on low mapping rates.

| Ecotype | Natal host | Treatment description | Treatment code | Sequenced | Passed quality filters |
|---|---|---|---|---|---|
| CH | *C. heterophyllum* (ancestral) | Control | H | 7 | 7 |
| | | Cross-foster to natal host | HH | 7 | 7 |
| | | Cross-foster to novel host | HO | 7 | 7 |
| CO | *C. oleraceum* (derived) | Control | O | 6 | 6 |
| | | Cross-foster to natal host | OO | 7 | 6 |
| | | Cross-foster to novel host | OH | 6 | 6 |

***Table S2. Sample details.***

Sample ID, ecotype, treatment, quality assessment of raw and trimmed reads, Salmon mapping rate and total mapped reads. [see SupplementaryTables.xlsx; dimensions: 40 x 17].

***Table S3. Gene set enrichment analysis of differentially expressed genes.***

GSEA on up and down regulated gene sets in each differential expression comparison (comp, e.g., H vs. O), where "down" genes are more expressed in the first treatment (e.g., H) and "up" genes are more expressed in the second treatment (e.g., O). Enrichment was tested using classic Fisher's exact tests, and Fisher's exact tests adjusted for GO-term dependency using the parent-child algorithm (TopGO; Alexa & Rahnenfuhrer, 2016). We use a parent-child p-value cutoff of 0.01 to determine whether a term was significantly enriched. We tested for enriched biological processes (BP) and molecular functions (MF). NAs represent sets for which no DE genes were detected. [see SupplementaryTables.xlsx; dimensions: 2857 x 11].

***Table S4. Gene set enrichment analysis of genes that were differentially expressed in both H vs. O and HH vs. OO comparisons.***

Enrichment was tested using classic Fisher's exact tests, and exact tests adjusted for GO-term dependency using the parent-child algorithm (TopGO; Alexa & Rahnenfuhrer, 2016). We use a parent-child p-value cutoff of 0.01 to determine whether a term was significantly enriched. We tested for enriched biological processes (BP) and molecular functions (MF). [see SupplementaryTables.xlsx; dimensions: 57 x 10].

***Table S5. Summary of weighted gene coexpression network modules.***

Membership was calculated as the correlation between expression and module eigengene of a given module. Module 'centers' for each sample were calculated as the mean expression of genes with 0.6 or greater membership in a given module. [see SupplementaryTables.xlsx; dimensions: 819 x 7].

***Table S6. Differentially expressed genes overlapping outlier windows.***

Gene functions and names were from the default EggNOG emapper functional annotation. Amino acid sequences of the longest isoform from each locus were also blasted against Drosophila melanogaster annotated proteins in Flybase.org. Here we report the name, expectation (e) value, and processes/functions of the highest Blastp match.

| **geneID** | **Overlaps** | **Gene function** | **Gene name** | **Flybase match** | **Blastp e-value** | **Biological process/molecular function** |
|---|---|---|---|---|---|---|
| Tcon_g11002 | Outlier, up in H | Sequence-specific DNA binding transcription factor activity | NFAT5 | Dmel\NFAT (CG11172) | < 1.0e-126 | Predicted to enable DNA-binding transcription factor activity, RNA polymerase II-specific and RNA polymerase II cis-regulatory region sequence-specific DNA binding activity. Involved in negative regulation of synaptic vesicle exocytosis and response to salt stress. Predicted to be part of the transcription regulator complex. |
| Tcon_g14432 | Outlier, up in H | Kinesin binding | KIFAP3 | Dmel\Kap3 (CG11759) | < 1.0e-126 | MF: kinesin binding; protein binding. BP: cellular process; sensory perception; nervous system process; microtubule-based movement; cellular component organization or biogenesis |
| Tcon_g16891 | Outlier, up in H | nucleic acid binding | - | Dmel\CG12877 | 2.7E-44 | MF: exonuclease activity; nucleic acid binding; RNA exonuclease-like domain |
| Tcon_g18210 | Outlier, up in H | 5' nucleotidase family | NT5C2 | Dmel\Nt5b (CG32549) | < 1.0e-126 | MF: 5'-nucleotidase activity. BP: adenosine metabolic process |
| Tcon_g23468 | Outlier, up in H | Partitioning defective 3 | PARD3 | Dmel\baz (CG5055) | < 1.0e-126 | Scaffold protein that forms a complex with the products of par-6 and aPKC and with other cortical, cytoskeletal and **regulatory proteins**. MF: phosphatidylinositol binding; protein binding; phosphatidic acid binding. BP 26 terms: protein localization; establishment of cell polarity; macromolecule localization; cellular localization; axis elongation |
| Tcon_g5492 | Outlier, up in H | ubiquitinyl hydrolase activity | USP2 | Dmel\Usp2 (CG14619) | < 1.0e-126 | MF: K48-linked deubiquitinase activity; proteasome binding; cysteine-type deubiquitinase activity. BP 7 |

| | | | | | | |
|---|---|---|---|---|---|---|
| | | | | | | terms: nitrogen compound metabolic process; post-translational protein modification; organic substance catabolic process; regulation of transport; negative regulation of antimicrobial peptide production |
| Tcon_g5788 | Outlier, up in H | histone-lysine N-methyltransferase activity | SETD2 | Dmel\Set2 (CG1716) | < 1.0e-126 | SET domain containing 2 (Set2) encodes an essential histone methyltransferase that marks active gene bodies with H3K36me3; MF 7 terms: transferase activity; histone H3 methyltransferase activity; methyltransferase activity; lysine N-methyltransferase activity; protein methyltransferase activity. BP: instar larval development; retrotransposon silencing by heterochromatin formation; wing disc development; regulation of DNA-templated transcription; ecdysone receptor-mediated signaling pathway |
| Tcon_g8783 | Outlier, up in H | otopetrin 3 | OTOP3 | Dmel\OtopLa (CG42492) | < 1.0e-126 | MF: sour taste receptor activity; proton channel activity. BP: detection of chemical stimulus involved in sensory perception of sour taste; proton transmembrane transport. |
| Tcon_g15631 | Outlier, up in HH | - | - | Dmel\CG1545 | 4.8E-40 | MF: unknown. BP: unknown |
| Tcon_g14351 | Outlier, up in HH | - | - | CG13012 | 2.4E-18 | BP: unknown |
| Tcon_g23712 | Outlier, up in H, up in HH | Belongs to the class-III pyridoxal-phosphate-dependent aminotransferase family | PHYKPL | Dmel\CG8745 | < 1.0e-126 | MF: ethanolamine-phosphate phospho-lyase activity; pyridoxal phosphate binding; transaminase activity. BP: response to nicotine |
| Tcon_g206 | Outlier, up in O | Eukaryotic protein of unknown function (DUF846) | TVP23B | Dmel\CG5021 | 3.1E-98 | BP: vesicle-mediated transport; protein secretion; Human ortholog TVP23B (trans-golgi network vesicle protein 23 homolog B), especially expressed in larval digestive system |
| Tcon_g5466 | Outlier, up in O | - | - | Dmel\tty (CG1693) | 1.97199 | MF: volume-sensitive chloride channel activity; intracellular calcium activated chloride channel |

| | | | | | | |
|---|---|---|---|---|---|---|
| | | | | | | activity; chloride channel activity. BP: chloride transport. |
| Tcon_g5479 | Outlier, up in O | Oxidoreductase activity. BP metabolic process | HSD17B11 | Dmel\Ldsdh1 (CG2254) | 1.7E-126 | MF: oxidoreductase activity, acting on the CH-OH group of donors, NAD or NADP as acceptor |
| Tcon_g11824 | Outlier, up in OO | Chitin-binding domain type 2 | Cpap3-d2 | Dmel\Gasp (CG10287) | 5.8E-21 | MF: chitin binding. BP: chitin-based cuticle development; regulation of tube size, open tracheal system. |
| Tcon_g5465 | Outlier, up in O, up in OO | PPIases accelerate the folding of proteins. It catalyzes the cis-trans isomerization of proline imidic peptide bonds in oligopeptides | Cyclophilin 1 | Dmel\Cyp1 (CG9916) | 3.5E-80 | MF: cyclosporin A binding; peptidyl-prolyl cis-trans isomerase activity. BP: protein folding; response to oxidative stress |

***Table S7. Kruskall-Wallis tests comparing population genomic metrics for H vs. O differentially expressed (DE) genes.***

DE and not DE genes were compared inside and outside of the inversion. Tests were performed on all not-DE genes, and also on equal sized sets of not-DE genes matched for gene length. P-values were corrected for multiple comparisons using the Benjamini-Hochberg method.

| Inversion | Metric | N | KW stat. | df | p-value | dataset | effsize | magnitude | Adj. p (BH) |
|---|---|---|---|---|---|---|---|---|---|
| Inside | Fst | 652 | 2.02 | 1 | $1.56x10^{-01}$ | all | $1.56x10^{-03}$ | small | 1.00 |
| Outside | Fst | 12531 | 3.50 | 1 | $6.12x10^{-02}$ | all | $2.00x10^{-04}$ | small | $9.20x10^{-01}$ |
| Inside | Fst | 104 | 3.61 | 1 | $5.75x10^{-02}$ | matched | $2.56x10^{-02}$ | small | $9.20x10^{-01}$ |
| Outside | Fst | 1590 | 7.02 | 1 | $8.06x10^{-03}$ | matched | $3.79x10^{-03}$ | small | $1.77x10^{-01}$ |
| Inside | dxy | 652 | 1.27 | 1 | $2.61x10^{-01}$ | all | $4.09x10^{-04}$ | small | 1.00 |
| Outside | dxy | 12528 | 34.57 | 1 | $4.11x10^{-09}$ | all | $2.68x10^{-03}$ | small | **$1.32x10^{-07}$** |
| Inside | dxy | 108 | 5.05 | 1 | $2.47x10^{-02}$ | matched | $3.82x10^{-02}$ | small | $4.20x10^{-01}$ |
| Outside | dxy | 1586 | 31.77 | 1 | $1.74x10^{-08}$ | matched | $1.94x10^{-02}$ | small | **$5.39x10^{-07}$** |
| Inside | Δπ | 644 | 2.82 | 1 | $9.32x10^{-02}$ | all | $2.83x10^{-03}$ | small | 1.00 |
| Outside | Δπ | 12257 | 1.23 | 1 | $2.67x10^{-01}$ | all | $1.88x10^{-05}$ | small | 1.00 |
| Inside | Δπ | 112 | 3.41 | 1 | $6.47x10^{-02}$ | matched | $2.19x10^{-02}$ | small | $9.20x10^{-01}$ |
| Outside | Δπ | 1562 | 3.35 | 1 | $6.74x10^{-02}$ | matched | $1.50x10^{-03}$ | small | $9.20x10^{-01}$ |
| Inside | ΔD | 644 | 6.00 | 1 | $1.43x10^{-02}$ | all | $7.79x10^{-03}$ | small | $2.57x10^{-01}$ |
| Outside | ΔD | 12257 | 1.12 | 1 | $2.91x10^{-01}$ | all | $9.49x10^{-06}$ | small | 1.00 |
| Inside | ΔD | 112 | 6.88 | 1 | $8.73x10^{-03}$ | matched | $5.34x10^{-02}$ | small | $1.77x10^{-01}$ |
| Outside | ΔD | 1562 | 0.02 | 1 | $8.82x10^{-01}$ | matched | $-6.27x10^{-04}$ | small | 1.00 |
| Inside | π (CH) | 791 | 7.61 | 1 | $5.81x10^{-03}$ | all | $8.37x10^{-03}$ | small | $1.39x10^{-01}$ |
| Outside | π (CH) | 12444 | 27.12 | 1 | $1.91x10^{-07}$ | all | $2.10x10^{-03}$ | small | **$5.54x10^{-06}$** |
| Inside | π (CH) | 142 | 11.65 | 1 | $6.43x10^{-04}$ | matched | $7.61x10^{-02}$ | moderate | **$1.67x10^{-02}$** |
| Outside | π (CH) | 1584 | 25.16 | 1 | $5.27x10^{-07}$ | matched | $1.53x10^{-02}$ | small | **$1.48x10^{-05}$** |
| Inside | π (CO) | 664 | 9.56 | 1 | $1.99x10^{-03}$ | all | $1.29x10^{-02}$ | small | **$4.98x10^{-02}$** |
| Outside | π (CO) | 12956 | 29.59 | 1 | $5.33x10^{-08}$ | all | $2.21x10^{-03}$ | small | **$1.60x10^{-06}$** |
| Inside | π (CO) | 110 | 7.12 | 1 | $7.62x10^{-03}$ | matched | $5.67x10^{-02}$ | small | $1.75x10^{-01}$ |
| Outside | π (CO) | 1614 | 21.71 | 1 | $3.17x10^{-06}$ | matched | $1.28x10^{-02}$ | small | **$8.56x10^{-05}$** |
| Inside | Taj. D (CH) | 791 | 0.42 | 1 | $5.16x10^{-01}$ | all | $-7.34x10^{-04}$ | small | 1.00 |
| Outside | Taj. D (CH) | 12444 | 1.12 | 1 | $2.90x10^{-01}$ | all | $9.80x10^{-06}$ | small | 1.00 |
| Inside | Taj. D (CH) | 142 | 0.02 | 1 | $8.78x10^{-01}$ | matched | $-6.98x10^{-03}$ | small | 1.00 |
| Outside | Taj. D (CH) | 1584 | 2.64 | 1 | $1.04x10^{-01}$ | matched | $1.04x10^{-03}$ | small | 1.00 |
| Inside | Taj. D (CO) | 664 | 6.99 | 1 | $8.21x10^{-03}$ | all | $9.04x10^{-03}$ | small | $1.77x10^{-01}$ |
| Outside | Taj. D (CO) | 12956 | 0.37 | 1 | $5.43x10^{-01}$ | all | $-4.86x10^{-05}$ | small | 1.00 |
| Inside | Taj. D (CO) | 110 | 6.65 | 1 | $9.93x10^{-03}$ | matched | $5.23x10^{-02}$ | small | $1.89x10^{-01}$ |
| Outside | Taj. D (CO) | 1614 | 0.00 | 1 | $9.71x10^{-01}$ | matched | $-6.20x10^{-04}$ | small | 1.00 |

***Table S8. Kruskall-Wallis tests comparing population genomic metrics for HH vs. OO DE genes.***

Genes that were DE and not-DE were compared inside and outside of the inversion. Tests were performed on all not-DE genes, and also on equally-sized sets of not-DE genes matched for gene length. P-values were corrected for multiple comparisons using the Benjamini-Hochberg method.

| Inversion | Metric | N | KW stat. | df | p-value | dataset | effsize | magnitude | Adj. p (BH) |
|---|---|---|---|---|---|---|---|---|---|
| Inside | Fst | 652 | 3.394 | 1 | 0.065 | all | $3.68x10^{-03}$ | small | 1.000 |
| Outside | Fst | 12531 | 0.002 | 1 | 0.964 | all | $-7.97x10^{-05}$ | small | 1.000 |
| Inside | Fst | 32 | 0.853 | 1 | 0.356 | matched | $-4.91x10^{-03}$ | small | 1.000 |
| Outside | Fst | 90 | 0.069 | 1 | 0.793 | matched | $-1.06x10^{-02}$ | small | 1.000 |
| Inside | dxy | 652 | 0.082 | 1 | 0.775 | all | $-1.41x10^{-03}$ | small | 1.000 |
| Outside | dxy | 12528 | 0.261 | 1 | 0.609 | all | $-5.90x10^{-05}$ | small | 1.000 |
| Inside | dxy | 32 | 0.043 | 1 | 0.836 | matched | $-3.19x10^{-02}$ | small | 1.000 |
| Outside | dxy | 90 | 0.055 | 1 | 0.815 | matched | $-1.07x10^{-02}$ | small | 1.000 |
| Inside | $\Delta\pi$ | 644 | 3.818 | 1 | 0.051 | all | $4.39x10^{-03}$ | small | 1.000 |
| Outside | $\Delta\pi$ | 12257 | 0.055 | 1 | 0.814 | all | $-7.71x10^{-05}$ | small | 1.000 |
| Inside | $\Delta\pi$ | 32 | 0.998 | 1 | 0.318 | matched | $-7.07x10^{-05}$ | small | 1.000 |
| Outside | $\Delta\pi$ | 86 | 0.911 | 1 | 0.340 | matched | $-1.06x10^{-03}$ | small | 1.000 |
| Inside | $\Delta D$ | 644 | 4.391 | 1 | 0.036 | all | $5.28x10^{-03}$ | small | 0.975 |
| Outside | $\Delta D$ | 12257 | 0.079 | 1 | 0.779 | all | $-7.52x10^{-05}$ | small | 1.000 |
| Inside | $\Delta D$ | 32 | 1.075 | 1 | 0.300 | matched | $2.49x10^{-03}$ | small | 1.000 |
| Outside | $\Delta D$ | 86 | 0.471 | 1 | 0.492 | matched | $-6.29x10^{-03}$ | small | 1.000 |
| Inside | $\pi$ (CH) | 791 | 4.508 | 1 | 0.034 | all | $4.45x10^{-03}$ | small | 0.944 |
| Outside | $\pi$ (CH) | 12444 | 1.288 | 1 | 0.256 | all | $2.32x10^{-05}$ | small | 1.000 |
| Inside | $\pi$ (CH) | 44 | 1.378 | 1 | 0.240 | matched | $8.99x10^{-03}$ | small | 1.000 |
| Outside | $\pi$ (CH) | 88 | 1.037 | 1 | 0.309 | matched | $4.25x10^{-04}$ | small | 1.000 |
| Inside | $\pi$ (CO) | 664 | 5.379 | 1 | 0.020 | all | $6.61x10^{-03}$ | small | 0.592 |
| Outside | $\pi$ (CO) | 12956 | 0.728 | 1 | 0.394 | all | $-2.10x10^{-05}$ | small | 1.000 |
| Inside | $\pi$ (CO) | 32 | 6.189 | 1 | 0.013 | matched | $1.73x10^{-01}$ | large | 0.400 |
| Outside | $\pi$ (CO) | 88 | 0.263 | 1 | 0.608 | matched | $-8.56x10^{-03}$ | small | 1.000 |
| Inside | Taj. D (CH) | 791 | 0.900 | 1 | 0.343 | all | $-1.27x10^{-04}$ | small | 1.000 |
| Outside | Taj. D (CH) | 12444 | 0.654 | 1 | 0.419 | all | $-2.78x10^{-05}$ | small | 1.000 |
| Inside | Taj. D (CH) | 44 | 0.014 | 1 | 0.907 | matched | $-2.35x10^{-02}$ | small | 1.000 |
| Outside | Taj. D (CH) | 88 | 0.361 | 1 | 0.548 | matched | $-7.43x10^{-03}$ | small | 1.000 |
| Inside | Taj. D (CO) | 664 | 5.577 | 1 | 0.018 | all | $6.91x10^{-03}$ | small | 0.546 |
| Outside | Taj. D (CO) | 12956 | 0.225 | 1 | 0.635 | all | $-5.98x10^{-05}$ | small | 1.000 |
| Inside | Taj. D (CO) | 32 | 6.962 | 1 | 0.008 | matched | $1.99x10^{-01}$ | large | 0.267 |
| Outside | Taj. D (CO) | 88 | 0.215 | 1 | 0.643 | matched | $-9.13x10^{-03}$ | small | 1.000 |

***Table S9. Dunn's tests comparing population genomic metrics among weighted gene coexpression network modules.***

P-values were corrected for multiple comparisons within each population genomic metric (n = 190 tests) using the Benjamini-Hochberg method. [see SupplementaryTables.xlsx; dimensions: 1520 x 9].

### Supplementary Figures

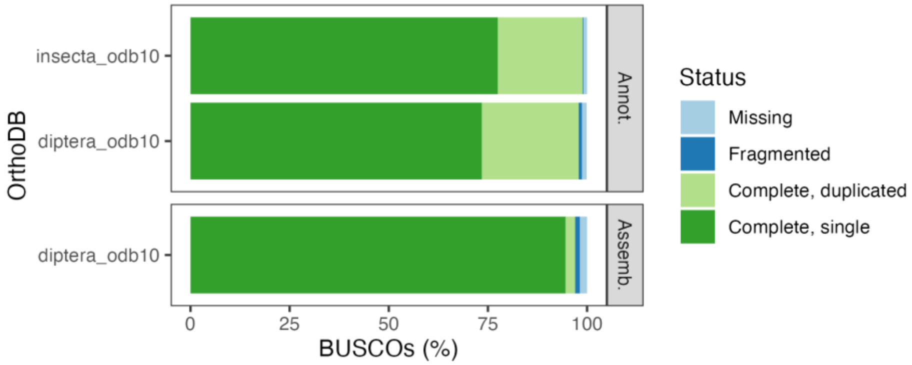

***Figure S1. BRAKER3 annotation quality.***

Completeness of the amino acid sequences was assessed using Insecta and Diptera single copy ortholog databases (OrthoDB v. 10). Overall completeness (single + duplicated orthologs) of the annotation was similar to that of the genome assembly.

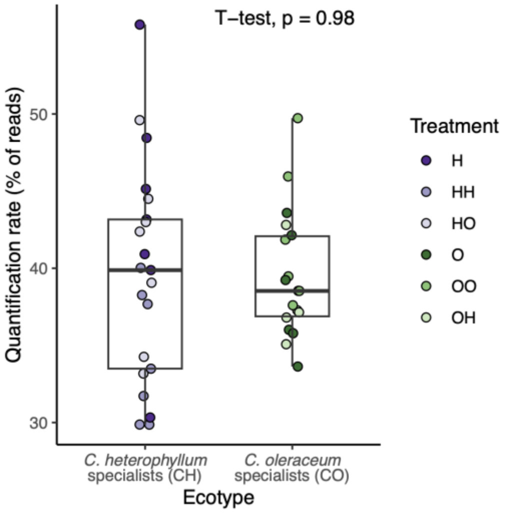

***Fig. S2. Quantification rates of trimmed reads aligned to BRAKER3 transcripts using Salmon.***

There was no difference in quantification rates between the two ecotypes, confirming no overall mapping bias resulting from mapping to a genome generated from a male *C. heterophyllum* specialist. Box plots show 25th, 50th, and 75th quartiles. Whiskers extend no more than 1.5 times the interquartile range.

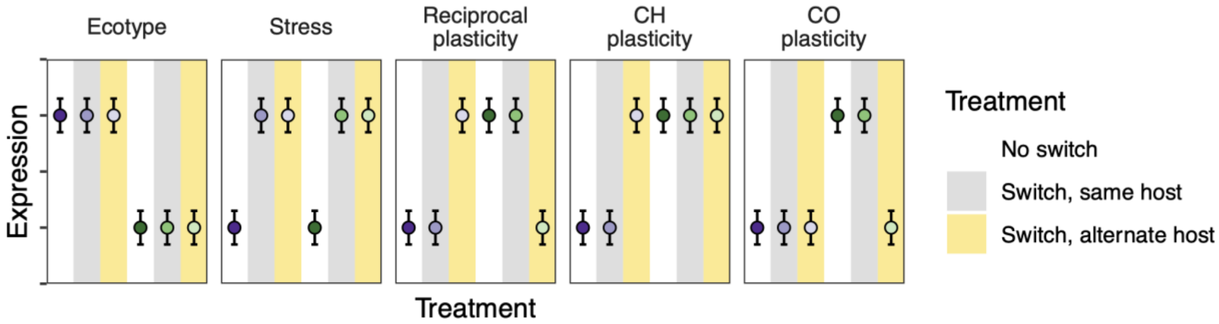

***Figure S3. Predicted patterns under five scenarios of gene expression among cross-fostered larvae.***

(1) Expression correlates with ecotype, consistently higher in one ecotype than another, regardless of the host plant on which they are feeding. (2) Expression correlates with stress, or more specifically, larvae that have undergone the cross-fostering treatment compare to control larvae. (3) Expression shows reciprocal plasticity, in which expression in cross-fostered larvae reflect the host plant on which they are feeding. (4 and 5) Expression shows ecotype specific plasticity, where only larvae from one ecotype (CH or CO) shift their expression to reflect the host plant on which they are feeding. Background colors indicate whether larvae were taken directly from hosts (white), switched to the natal host (grey) or switched to the novel host (yellow).

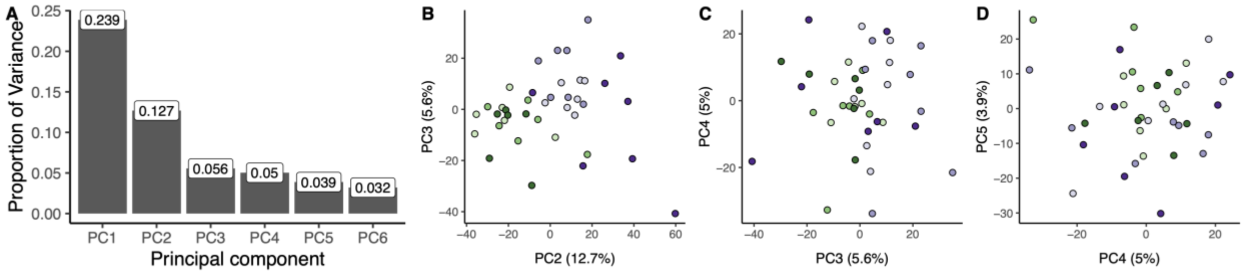

***Figure S4. Principal component analysis of normalized, transformed expression.***

(A) Proportion of the variance explained by the first six PC axes in analysis of the 500 genes with the most variable expression. Samples tended to cluster by ecotype on the second PC axis (B), but not on axes 3, 4, or 5 (C, D).

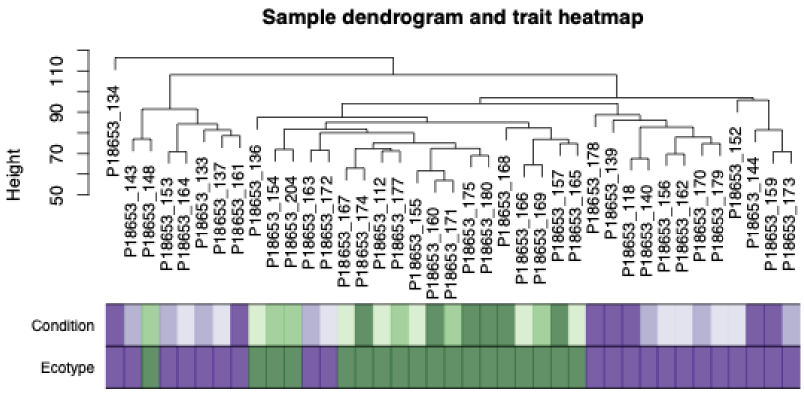

***Figure S5. Hierarchical clustering of larval samples based on regularized log-transformed and normalized gene expression.***

The count matrix was filtered to exclude genes with < 5 reads in ≤ 5 samples. Filtered read counts were normalized and transformed (regularized log) before clustering. CO larvae (ecotype = dark green) tended to cluster together. There was little clustering within ecotype by treatment/condition, i.e., harvested directly from host (dark purple or green), switched to the same host (medium purple or green), or switched to the alternative host (light purple or green).

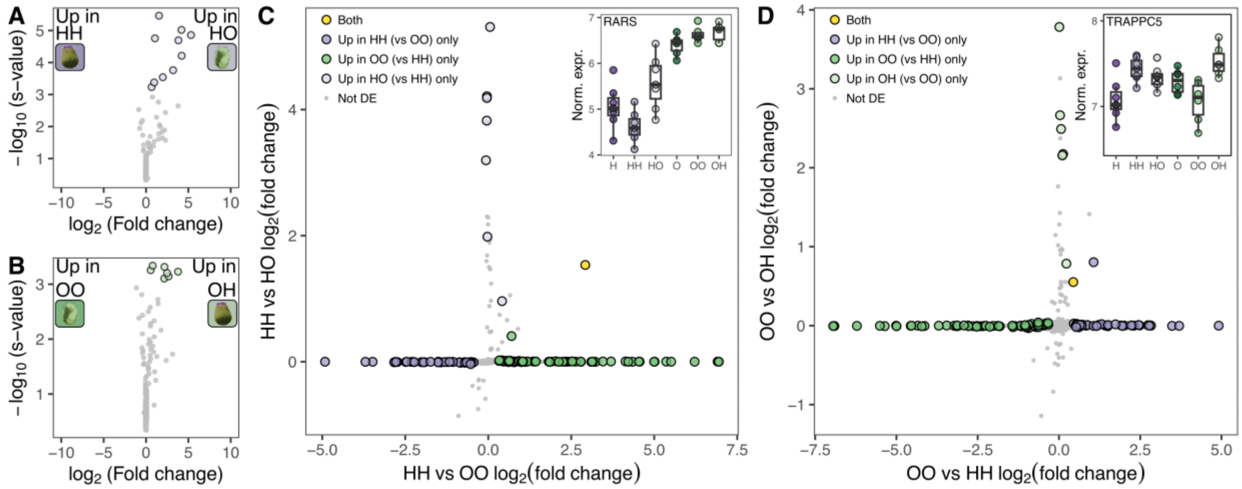

***Figure S6. Differential expression and expression plasticity in larvae cross-fostered to their natal host or a novel host.***

(A) Differentially expressed genes between CH larvae cross-fostered from *C. heterophyllum* to the same host (HH) and CH larvae cross-fostered from *C. heterophyllum* to *C. oleraceum* (HO). (B) Differentially expressed genes between OO and OH larvae. (C) Adaptive transcriptional plasticity in CH larvae was identified when genes that were differentially expressed between CH and CO larvae cross-fostered to their natal hosts (HH vs. OO) were also differentially expressed in CH larvae cross-fostered to *C. oleraceum* (HH vs. HO; yellow points). (D) Transcriptional plasticity in CO larvae cross-fostered to *C. heterophyllum*. Insets show normalized, regularized log-transformed expression of plastically expressed genes, (C) RARS and (D) TRAPPC5.

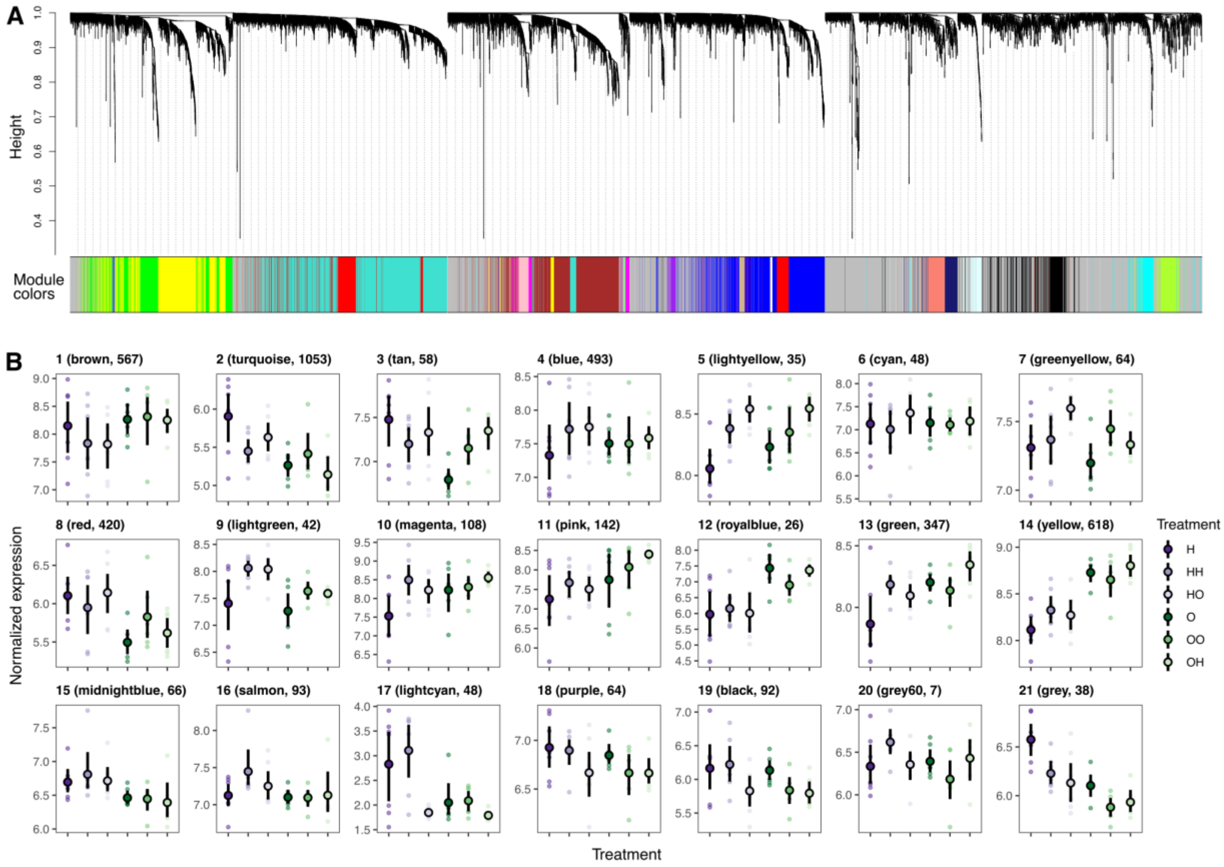

***Figure S7. Weighted gene co-expression module clustering and normalized expression across treatments.***

(A) Genes clustered into 21 modules using signed biweight midcorrelations. Genes were clustered using adjacency. (B) Transformed and normalized expression (regularized log transformation) of module centers differed among modules and treatments. Modules are numbered, with assigned colors and number of genes with membership >0.6 in parentheses. Small points show module centers for each sample, and black points and vertical lines represent means and 95% confidence intervals, respectively.

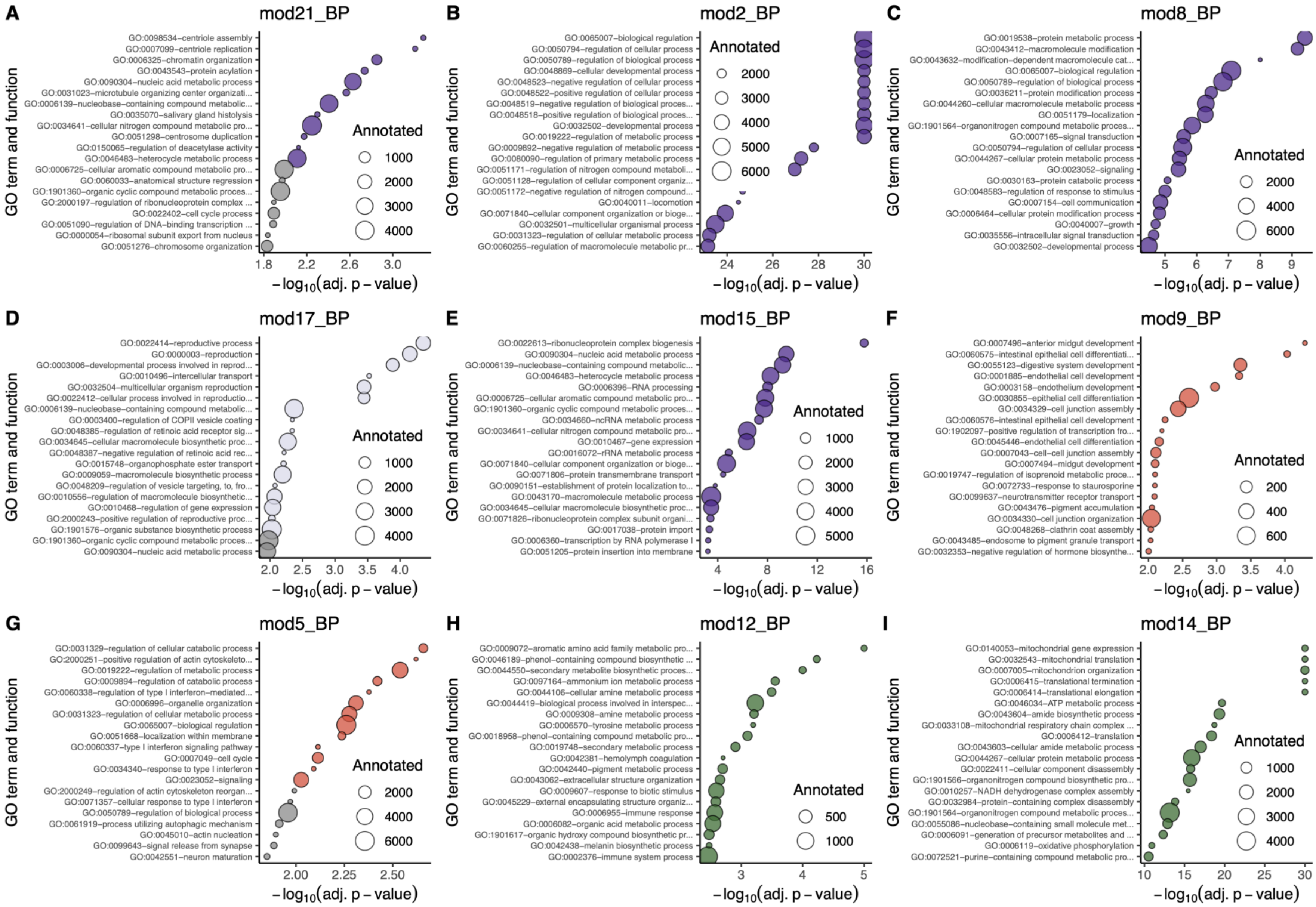

***Figure S8. Enriched biological processes in modules correlated with ecotype (21, 2, 8, 17, 15, 12, 14), stress (9, 5) or CH plasticity (21, 17, 14).*** Top 20 significantly enriched terms ($p < 0.01$) are colored according to the predictor with which they were most-highly correlated (Figure 4A): ecotype (up in CH = purple, up in CO = green), stress (up in cross-fostered = red), and CH plasticity (up in H and HH = light purple). Point size is scaled to reflect the number genes annotated with a given term in the functional annotation.

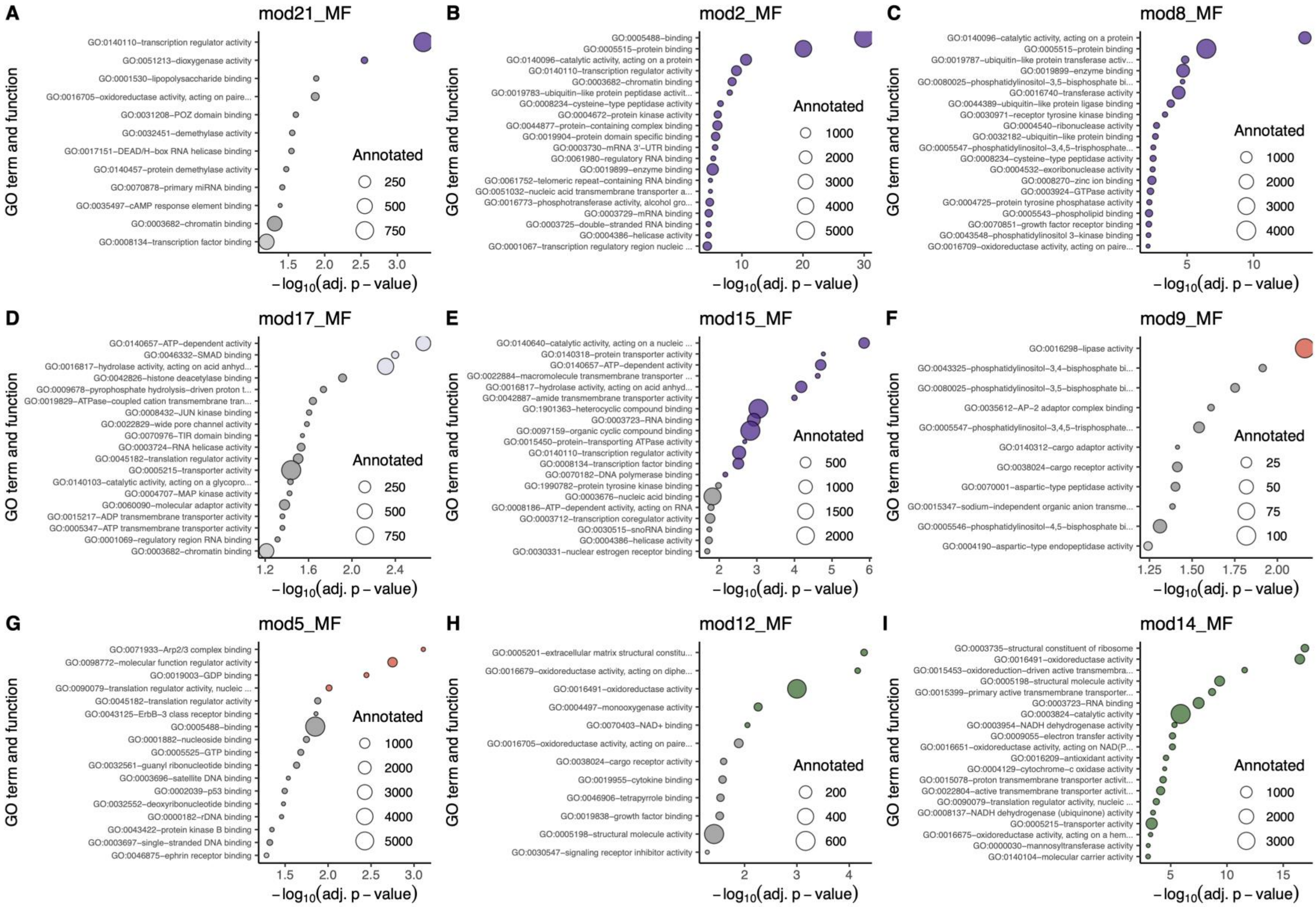

***Figure S9. Enriched molecular functions in modules correlated with ecotype (21, 2, 8, 17, 15, 12, 14), stress (9, 5) or CH plasticity (21, 17, 14).*** Top significantly enriched terms ($p < 0.01$, max. 20 pictured) are colored according to the predictor with which they were most-highly correlated (Figure 4A): ecotype (up in CH = purple, up in CO = green), stress (up in cross-fostered = red), and CH plasticity (up in H and HH = light purple). Point size is scaled to reflect the number genes annotated with a given term in the functional annotation.

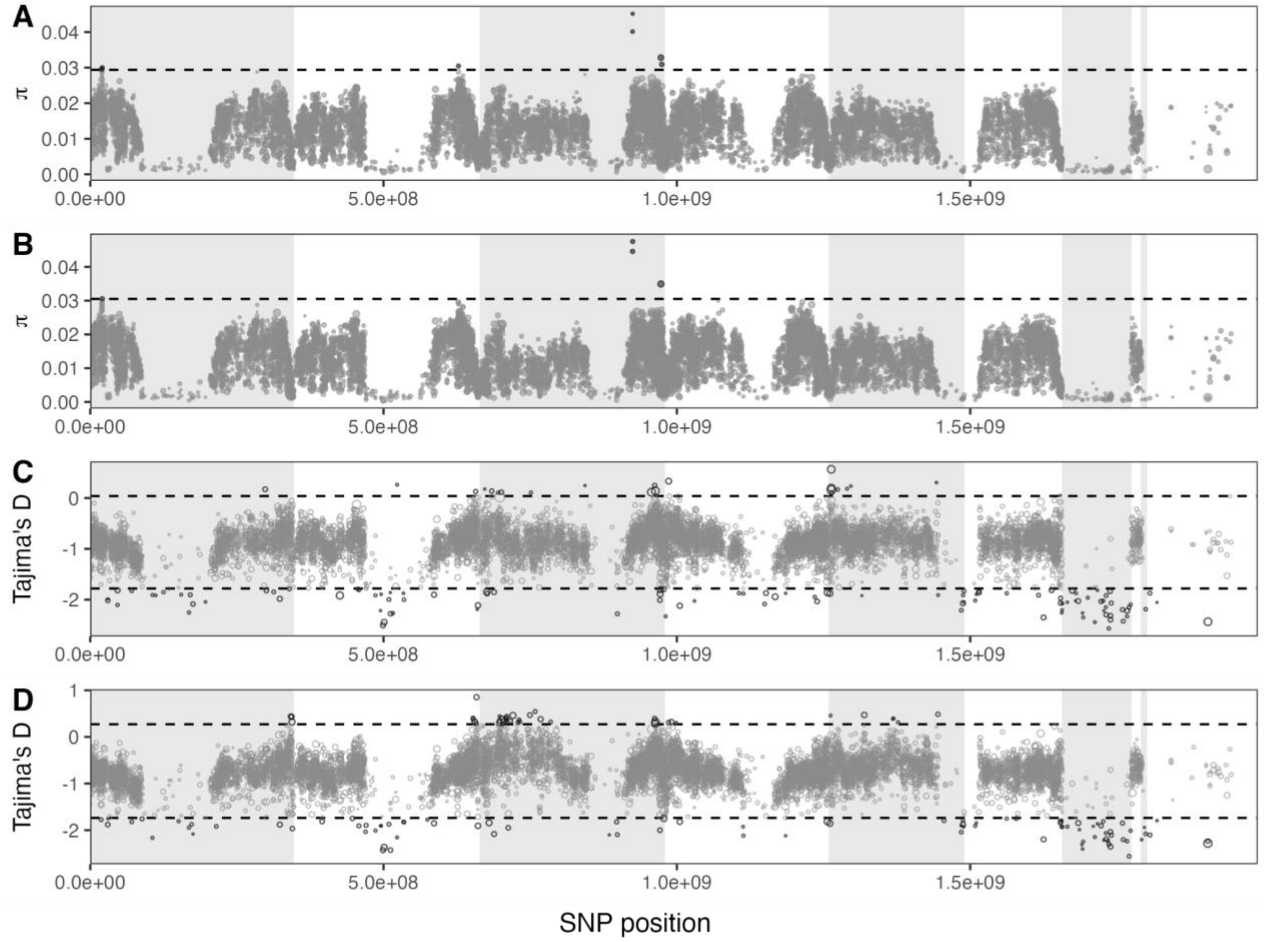

***Figure S10. Nucleotide diversity (π) and Tajima's D in CH and CO populations calculated over 50kb windows.***

Windows with fewer than 20% coverage were excluded, resulting in several gaps along the ordered contigs. Horizontal dashed lines represent the mean +/- 3 standard deviations, and outliers are colored in black. Contigs are ordered according to hypothetical linkage groups. Linkage groups are delineated as light gray or white bands, with the rightmost white band containing unscaffolded, low coverage contigs.

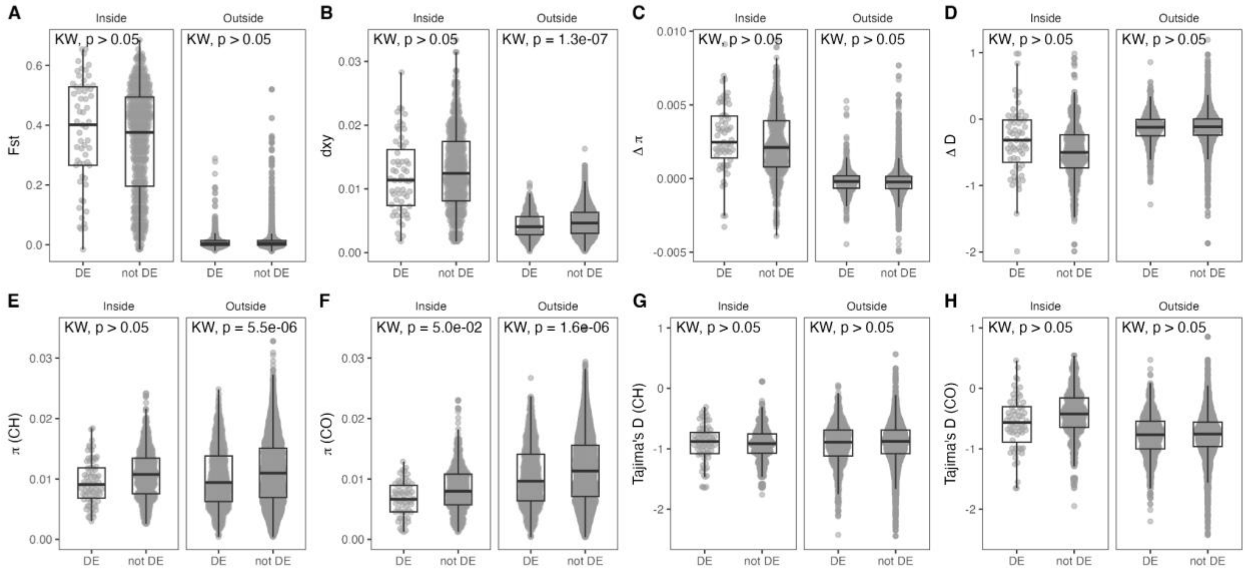

***Figure S11. Population genomic metrics for genes inside and outside of the putative inversion that were or were not differentially expressed (DE) between H and O larvae.***

Kruskal-Wallis tests were used to compare DE and not DE gene sets and p-values were corrected for 16 tests. Box plots show 25th, 50th, and 75th quartiles. Whiskers extend no more than 1.5 times the interquartile range. Outliers are not shown.

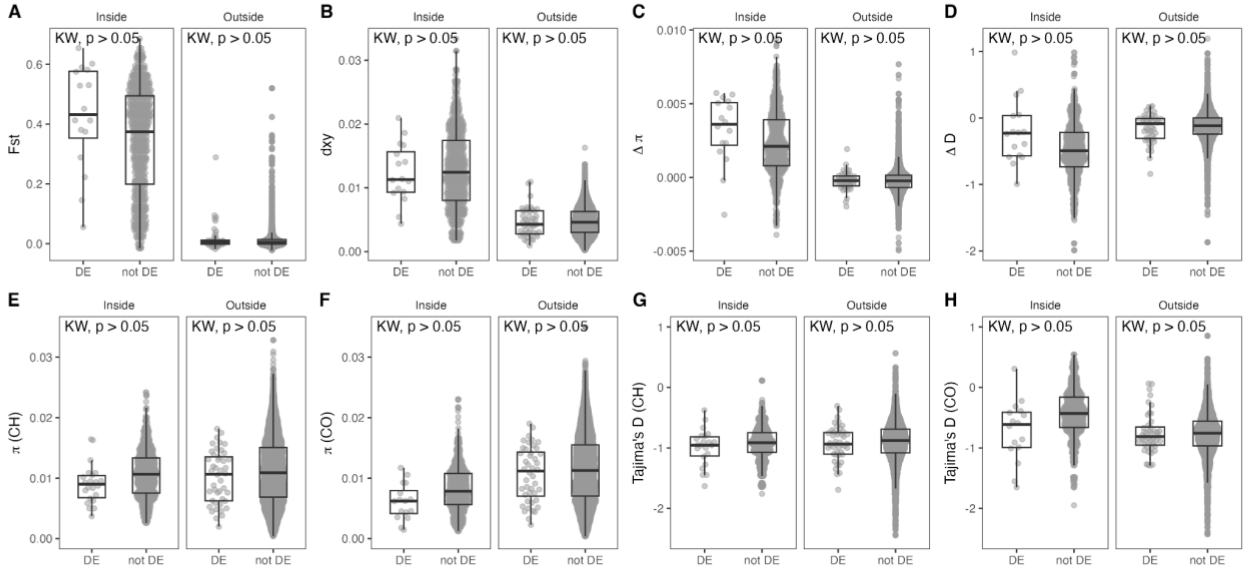

***Figure S12. Population genomic metrics for genes inside and outside of the putative inversion that were or were not differentially expressed (DE) between HH and OO larvae.***

Kruskal-Wallis tests were used to compare DE and not DE gene sets and p-values were corrected for 16 tests. Box plots show 25th, 50th, and 75th quartiles. Whiskers extend no more than 1.5 times the interquartile range. Outliers are not shown.

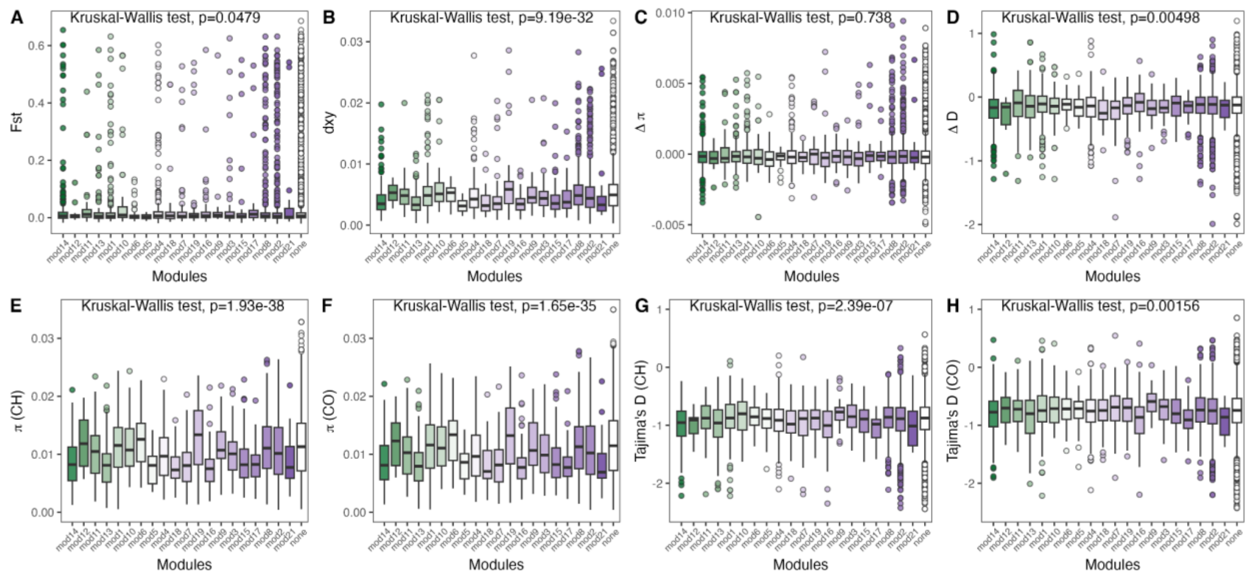

***Figure S13. Population genomic metrics among weighted gene coexpression modules***

Kruskal-Wallis tests were used to compare metrics across all modules, including genes that were not assigned to a module or did not meet the membership criteria (0.6) criteria to be retained in a module ("none"). P-values are corrected for 8 tests using the Benjamini-Hochberg correction. Dunn's tests were used to compare metrics between specific modules (Table S7). Box plots show 25th, 50th, and 75th quartiles. Whiskers extend no more than 1.5 times the interquartile range. Outliers beyond this range are shown as points. Boxes and points are colored according to the correlation of the module with ecotype (Figure 4A; "none" not colored).
